# Predicting the natural yeast phenotypic landscape with machine learning

**DOI:** 10.1101/2024.10.17.618784

**Authors:** Sakshi Khaiwal, Matteo De Chiara, Benjamin P Barré, Inigo Barrio-Hernandez, Simon Stenberg, Pedro Beltrao, Jonas Warringer, Gianni Liti

## Abstract

Most organisms’ traits result from the complex interplay of many genetic and environmental factors, making their prediction from genotypes difficult. Here, we used machine learning models to explore genotype-phenotype connections for 223 life history traits measured across 1011 genome-sequenced *Saccharomyces cerevisiae* strains. Firstly, we used genome-wide association studies to connect genetic variants with the phenotypes. Next, we benchmarked an automated machine learning pipeline that includes preprocessing, feature selection, and hyperparameters optimization in combination with multiple linear and complex machine learning methods. We determined gradient boosting machines as best performing in 65% of predictions and pangenome as best predictor, suggesting a considerable contribution of the accessory genome in controlling phenotypes. The accuracy broadly varied among the phenotypes (r = 0.2-0.9), consistent with varying levels of complexity, with stress resistance being easier to predict compared to growth across carbon and nitrogen nutrients. While no specific genomic features could be linked to the predictions for most phenotypes, machine learning identifies high-impact variants with established relationships to phenotypes despite being rare in the population. Near-perfect accuracies (r>0.95) were achieved when other phenomics data were used to aid predictions, suggesting shared useful information can be conveyed across phenotypes. Overall, our study underscores the power of machine learning to interpret the functional outcome of genetic variants.

## Main

The majority of traits on all biological levels are complex and controlled by multiple genetic loci, the environment, and interactions between the two. This interplay of many variables makes it difficult to determine their underlying genetic architecture^1,2^. Genome-wide association studies (GWAS) have proven successful in associating genetic markers to traits in species for which large cohorts can be sequenced and phenotyped^3–6^. However small effects and rare variants typically fail to pass the stringent statistical thresholds. For combinations of variants, the problem becomes exponentially bigger and only the strongest of common pairs can be detected. Therefore, we only have a partial view of the genetic architecture even for well studied traits^7^.

Machine learning (ML) is emerging as a powerful tool capable of constructing models that potentially can capture even complex genotype-phenotype relationships to enable predictions^8^ by identifying and prioritising causative variants^9^. These models can be further powered by implementing additional information, such as multi-omics^10^ and evolutionary frameworks^11^. However, both GWAS and predictions often lose power when environmental factors cannot be controlled^8^. Model organisms such as *Saccharomyces cerevisiae* are powerful systems for investigating these genotype-phenotype relationships due to the relative ease with which accurate genotypic and phenotypic information in controlled environments can be obtained^12–15^. The genome sequenced 1,011 *S. cerevisiae* collection, isolated worldwide from a broad variety of ecological sources, provided an accurate population-level variation portrait^16^. An ensemble of 223 life history phenotypes measured under controlled environmental conditions captures core components of the species’ life cycle such as growth, sporulation, survival under starvation conditions, and cell characteristics such as cell size, DNA content, and mitochondrial activity^16–19^. Both proteome and transcriptome of the 1,011 yeast collection have also been quantified, providing intermediate molecular traits that can aid predictions of higher-level phenotypes from genetic variants^20,21^. Furthermore, investigation of yeast natural variation can be coupled with multiplexed genome editing approaches, providing a platform to test ML predictions at scale^22,23^. However, an in-depth investigation of whether ML enables higher resolution of the yeast genotype-to-phenotype is currently lacking. Here, we systematically explore ML phenotypic predictions in the 1011 yeast collection across 223 phenotypes. To construct genotype-phenotype models that can enable predictions, we built a flexible ML pipeline to quantify the best methods and type of input information for predicting phenotypes and to build prediction models across the genotype and phenotype space of the collection.

## Results

### The yeast’s natural phenotypic landscape is shaped by correlations between unrelated traits

To define the phenotypic landscape of baker’s yeast, we compiled 190 published life history traits measured across the 1,011 *S. cerevisiae* collection in five main studies and complemented these with 33 additional traits (Supplementary Data 1-2). The aggregated phenome dataset comprises 223 phenotypes largely corresponding to yeast life cycle traits, including chronological life span (CLS), sporulation efficiency, and asexual reproduction (cell yield and doubling time) (Fig. 1a). Traits were measured across a range of controlled environments, including variations in type and degree of carbon and nitrogen sources availability and in exposure to drugs and abiotic stresses, such as temperature and salinity, and classified into 8 main groups, based on the type of trait or trait-environment pairs (Fig. 1a-b, Supplementary Data 3). We first explored patterns of co-variation between traits, through an all-vs.-all pairwise correlation analysis and found significant correlations to be both more abundant and stronger than anticorrelations (Pearson’s *r* > 0.5 for 6.5% of pairs vs. *r* < −0.3 for 0.1%) (Fig. 1c). Small, strongly correlated clusters are mostly driven by similar environments, or different concentrations of the same condition (Fig. 1c). Overall, trait type had a stronger impact than environment type on trait similarity, with e.g. cell yields in different environments grouped together (Fig. 1c). However, 13% of the correlations and 60% of anticorrelations emerged between phenotypes of different classes. For example, cell yield during growth in the presence of azoles, which block the formation of mature sterols^24^, positively correlated with CLS (Fig. 1c). Anticorrelations (*r* < −0.3) between cell yield and doubling, reflecting r/K trade-offs, were more common than for phenotype pairs (1.6% vs. 0.6%), but still rare, consistent with that such trade-off manifesting only under specific environments^25^.

**Fig. 1.**
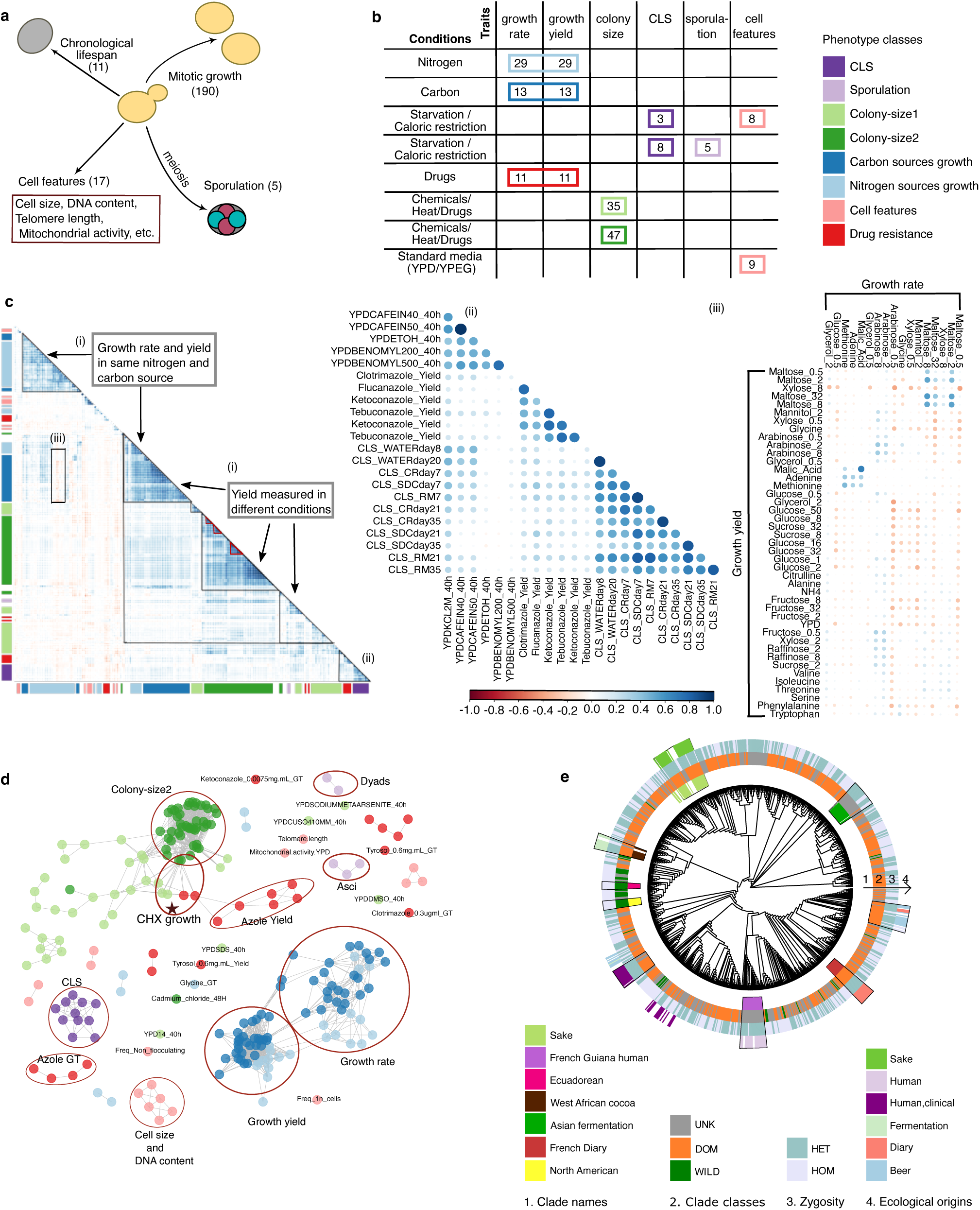
The phenotypic landscape. **a,** Schematic of measured phenotypes corresponding to key features of the yeast life cycle and cells. **b,** Phenotype classes based on trait and environmental conditions. **c,** Global phenotypic correlation map representing correlation/anticorrelation, strength and significance (respectively, dots colours, tones, and sizes). Colour bars on the *x* and *y* axes indicate phenotype classes. Small clusters show examples of strong correlations between phenotypes in similar conditions (red triangles). Larger correlation patterns (black lines) show (i) clustering of similar traits (yield) across environments, (ii) unexpected strong correlations between unrelated traits (CLS and drug resistance), (iii) trade-offs between growth rate and yield. **d,** Phenotypic networks based on correlation (>=0.5) reveal compact clusters within traits/environments and few disconnected phenotypes. Dots colour indicates phenotype classes. **e,** Phenomic tree of the 1011 *S. cerevisiae* circled by heat maps indicating clades, clade classes, heterozygosity and ecological origins. Clusters corresponding to clades are highlighted (black lines).

Next, we constructed phenotypic networks based on strongly correlated phenotypes (methods). Consistently, we observed major trait types, such as growth rate, yield, and CLS forming distinct subnetworks (Fig. 1d). The same or similar phenotypes measured in different studies were connected, as highlighted by the case of cycloheximide (CHX) (starred, Fig. 1d). Twelve phenotypes did not show strong correlation with other traits and remained unconnected (Fig. 1d). The low or no connectedness of these phenotypes is not due to higher experimental noise (Supplementary Data 4-5) and instead suggests they are controlled by distinct biological pathways. Networks of anti-correlations were substantially less frequent and weaker compared to correlations with most phenotypes being disconnected except for 6 small subnetworks (Supplementary Fig. 1a). Three of these subnetworks represent trade-offs between growth rate and yield in several sugars and drugs. Using the correlation matrix, we determined 4 to be the optimal number of clusters (based on silhouette scores) and hierarchical clustering with k=4 does recapitulate patterns observed in the correlation map (Supplementary Fig. 1b). For example, CLS clustered together with azole resistance, and growth yield across conditions from the three independent datasets clustered together.

To corroborate the above findings, we compared the natural phenotypic correlation patterns with those of single gene deletions derived from the reference laboratory strain background^26^. We retrieved 200 phenotypes measured in the deletion collection overlapping with our phenome dataset (Supplementary Data 6-7). Clustering analysis on the correlation matrix supported instances of shared correlation, including drug resistance and CLS (Supplementary Fig. 2a). Thus, these are likely true biological effects rather than spurious correlations due to shared bias.

Next, we investigated to what extent the co-variation in phenotypes agreed with overall genetic relatedness, and whether co-variation in specific phenotypes was driven by some specific yeast clades. Strains in a few clades, such as French Guiana, Far East Asia, and Asian fermentation were phenotypically similar (Fig. 1e). These clades showed very low intra-clade genetic variance, consistent with a higher degree of phenomic similarity (Supplementary Fig. 2b-d). Overall, comparing phenomic and genetic distances, either based on SNPs or gene presence-absence, we found no correlation. Thus, besides very closely related yeasts showing similar phenotypes, population structure, and phylogeny in general, appear to have next to no influence on *S. cerevisiae* intra-species phenotypic variation. Instead, wild and domesticated yeast clades clustered separately, despite different yeast clades having been domesticated independently at different time points and geographic regions. This is consistent with previous analysis^17^, and supports that domestication is the major determinant of yeast phenotypic variation.

### The yeast GWAS catalogue

To set a baseline for evaluating the capacity of machine-learning to explore the yeast genotype-phenotype map, we first established genotype-phenotype connections using current state-of-the-art GWAS approaches. We obtained a catalogue of 2341 SNPs, distributed across 1271 genes (Supplementary Data 8-10), associating to one or more phenotypes (Fig. 2a). Similarly, 34 loss-of-function variants (LOFs) in distinct genes, 65 gene presence/absence variations in the pangenome (PA) and 139 copy number variants (CNVs) associated with at least one trait. While other variants associated proportionally with all phenotype classes, CNVs were enriched in CLS and colony-size 1 and 2 (Supplementary Fig. 3a). Generally, the strength of phenotype associations, reflecting effect sizes, were lower for SNP than for other PA, CNVs and LOF variants (Supplementary Fig. 3b). The phenotype associated variants were mostly uniformly distributed across the genome, except for a strong enrichment associated with colony-size1 and 2 (from 34% genome-wide to 90%), which mostly reflects cell yield in stressful environments, within a chromosome XV region (Fig. 2a and Supplementary Fig. 3c). These variants mostly mapped to the stress response transcriptional activator *SFL1* and the adjacent actin-related chromatin remodeller *ARP8*^27^ (5 and 9 variants, associating with 18 and 20 phenotypes, respectively).

**Fig. 2.**
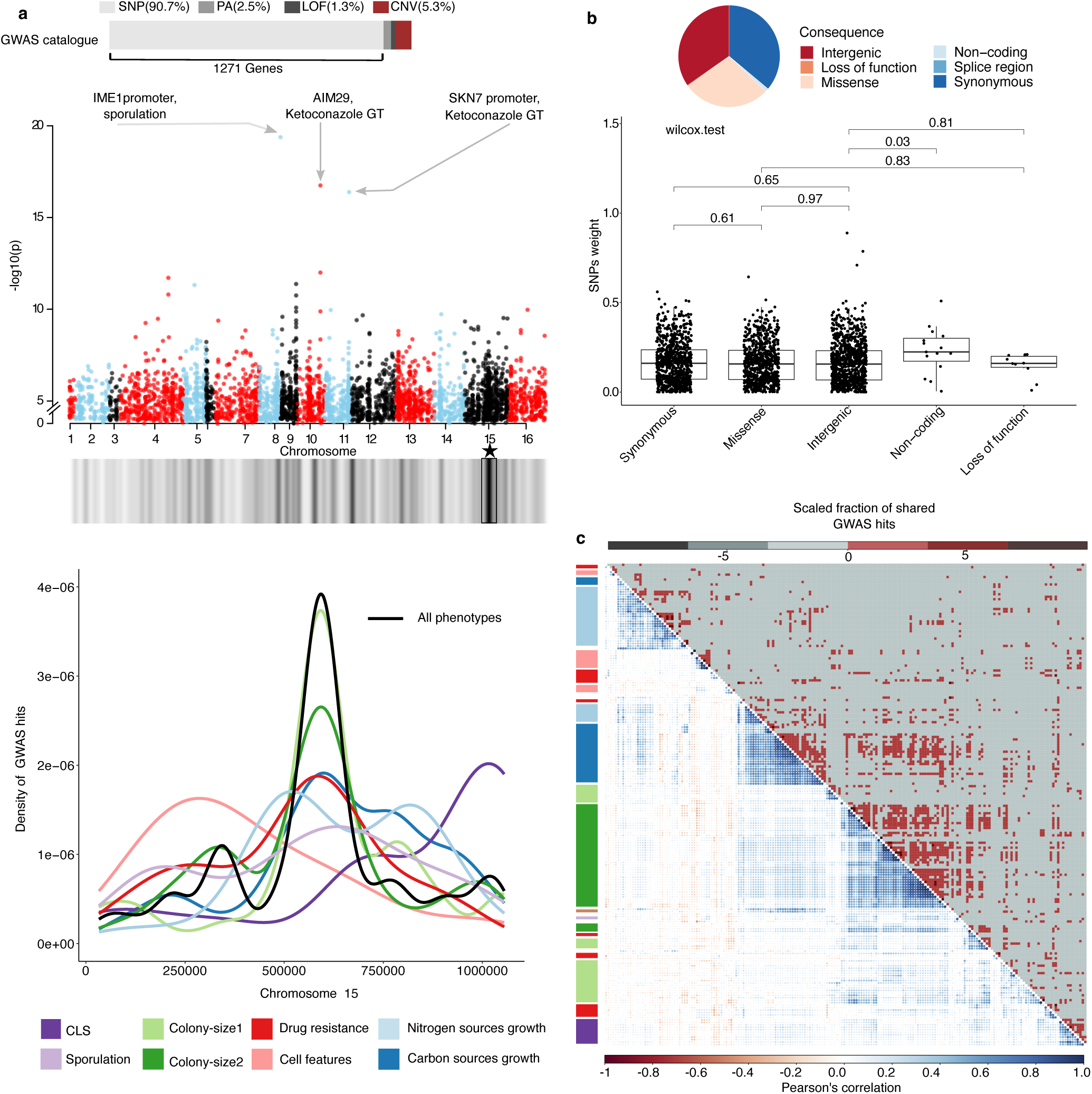
GWAS overview. **a,** Barplot (top) depicts the GWAS hits catalogue with SNPs constituting 90% of hits in 1271 genes, followed by CNVs (5.3%), PA (2.5), and LOF(1.3%). Genome-wide distribution of significant GWAS hits for the global phenome set. The region with the highest hit density (box) is zoomed-in (bottom panel) and lines colours indicate phenotype classes. **b**, Fractions of GWAS hits by type (pie chart) and weight distributions (box plot). **c**, Comparison between the phenomic correlation map (bottom left) and the scaled fraction of GWAS hits shared between phenotypes (top right).

Three SNPs had exceptionally high *p*-values (Fig. 2a) include a promoter variant (IME1_C-181T) in the master regulator *IME1* leading to sporulation defect and *AIM29* (c.159C>T, p.His53His) causing azole resistance, both of which we experimentally validated^17^ (*AIM29*, manuscript in preparation, https://github.com/SakshiKhaiwal/CEFIPRA), and *SKN7* promoter variant (SKN7_C-376G) within a highly conserved region associated to ketoconazole resistance. We used a variant effect predictor (VEP) to annotate all phenotype-associated variants. Surprisingly, synonymous (32%), missense (29%), and intergenic (37%) were equally represented among our GWAS hits (Fig. 2b), with fractions closely resembling how common these variant types are overall (Supplementary Fig. 3d). Furthermore, the strength of associations was similar for these variant categories (Fig. 2b). Using the mutfunc framework^28^, we found no enrichment of variants predicted to affect function or protein stability among phenotype associations, whereas protein interfaces had a low number of predictions available (Supplementary Data 11 and Supplementary Fig. 3e). Phenotype associated variants also affected essential and non-essential genes equally often (Supplementary Fig. 4a).

We calculated the fraction of GWAS hits that was shared between each pair of phenotypes (Fig. 2c) and found both correlated and anticorrelated phenotypes to share variants more often than expected by chance, validating that these relations have a genetic rather than an environmental (i.e. shared bias) basis (Supplementary Fig. 4b). Consistent with their strong correlations, cell yields across environments often shared GWAS hits (Fig. 2c), while the same variant sharing was not evident for correlated CLS and azole resistance phenotypes. Expanding this analysis to the gene-level by associating every polymorphic position to the corresponding gene (Supplementary Fig. 4c), we however found variants in the *FKS1*^29^ gene to associate with both CLS and azoles resistance, indicating defects in cell wall biosynthesis as a potential basis of their phenotypic correlation. We also identified three synonymous variants in the *SEC11* gene associated with the growth rate and yield trade-off in some sugar conditions, but swapping *SEC11* alleles between high and low-performing strains provided no conclusive support for causal effects (Supplementary Data 12-15), which could imply confounding intra or interlocus interactions.

Counting the number of phenotype associations per gene, we found 40% of the GWAS-associated genes (512 out of 1272) to be pleiotropic, with phenotypes reflecting cell-environment interactions, e.g. binding processes, cellular response to stimuli and signal transduction, being enriched among GWAS hits (Supplementary Data 16 and Supplementary Fig. 4d). Often, a few SNPs within these genes accounted for almost all of the pleiotropy, e.g. 2 SNPs in *CAC2* and 3 SNPs in *CDC23* associating with 28 and 24 phenotypes respectively, while a single SNP in the functionally uncharacterized YEL023C gene associated with 10 phenotypes.

We used a Personalized Page Rank (PPR) algorithm that tracks signal expansion through an interactome to identify pleiotropic gene groups^30,31^, using both our GWAS hits and published phenotype data on the yeast gene knockout collection as inputs. To identify pleiotropic gene modules that impact multiple complex traits, we extracted modules with more than 75% genes in common between the two traits (Supplementary Fig. 5a-b). For both natural and knockout collections, only a few such modules were identified between similar phenotypes within each collection individually, however, no overlap was found between similar phenotypes across the two collections (Supplementary Fig. 5a-b, Supplementary Data 17-18). For the natural collection, modules overlapping in multiple traits were mostly related to stress environments with enrichment for ubiquitin-like protein transferase activity and protein binding, while modules associated with sugar conditions were enriched in actin and cytoskeletal protein binding and other basal cell functions. However, few modules enriched for nucleic acid binding overlap between sugar conditions and stress environments (Supplementary Data 19-20).

### An integrated machine learning pipeline for yeast phenotype prediction

The availability of 223 phenotypes coupled with genetic features (SNPs, LOF, PA, CNVs, gene function disruption scores (P(AF) scores), Methods), and molecular-level (transcriptomic and proteomic) variations enabled us to predict yeast phenotypes. We constructed an all-in-one flexible pipeline (Gen-phen) for data pre-processing, feature selection and model learning to automate phenotype predictions (Methods and Fig. 3a). The preprocessing removes strains with missing phenotype values and features with too many unknown values (>25% of the strains), randomly splits the 1,011 *S. cerevisiae* strains into training (75%) and testing (25%) sets, transforms features on a standard scale (normal distribution) and finally infers missing feature values through imputation. For feature selection, the pipeline uses Lasso regression to select the features that best capture variation between strains and construct a sparse model that is computationally more efficient and generalizable for prediction.

**Figure 3.**
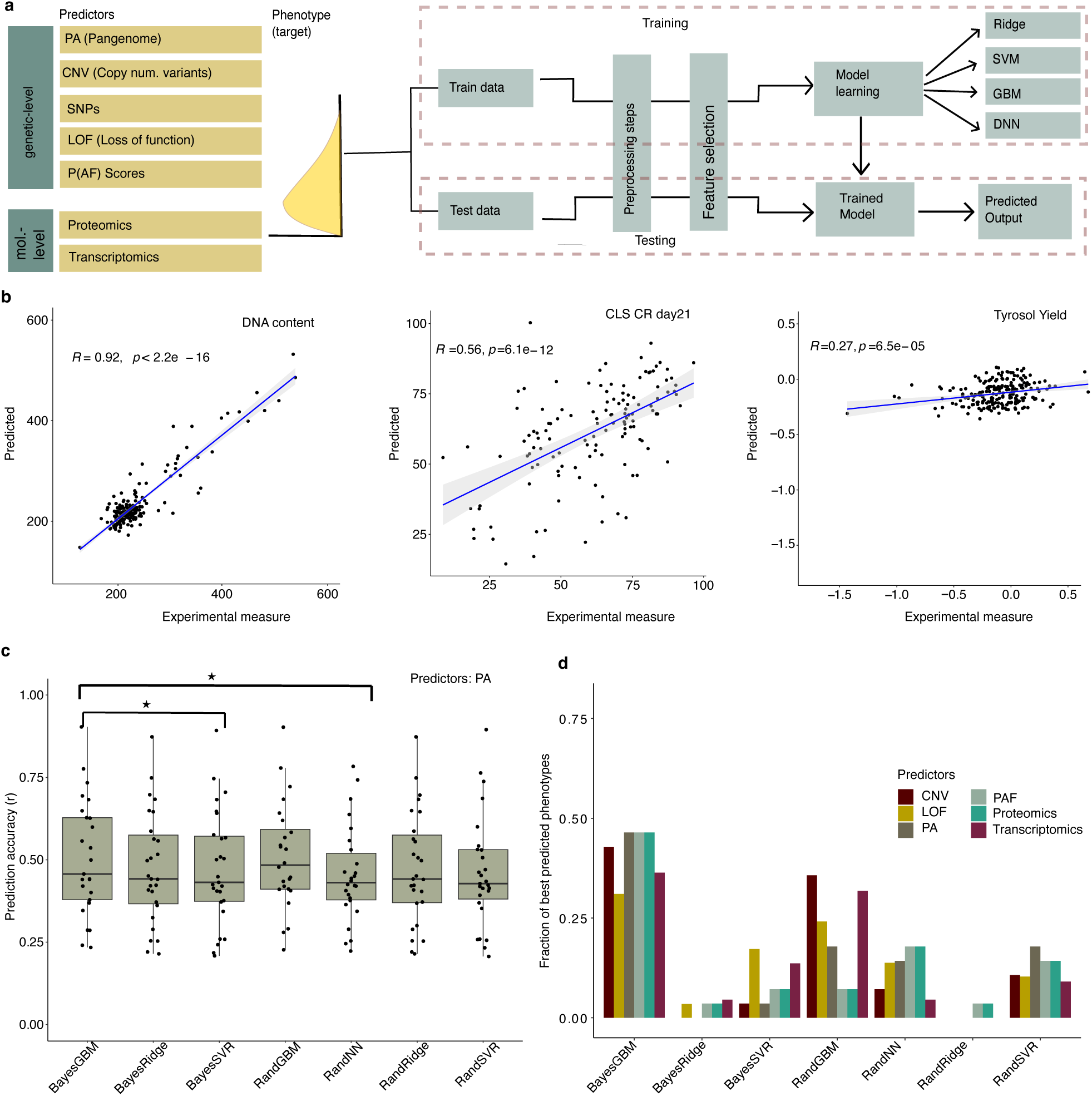
Benchmarking of the Gen-phen pipeline. **a,** Overview of the Gen-phen architecture. **b**, Examples of good (DNA content), moderate (CLS CR day21), and (Tyrosol yield) bad phenotypic predictions. **c,** Gen-phen benchmarking using 30 phenotypes shows no significant differences across methods except for the pairs Bayesian GBM - SVR, and Bayesian GBM - random NN (paired Wilcox test, p-value <=0.05). Only correlation coefficients with BF-corrected p-values less than 0.05 are shown. **d**, Fraction of the best performance for each method across predictors. Bayesian GBM was the best method (50%), followed by Random GBM and SVR.

In the model learning step, the user can choose between common machine learning algorithms (ridge, gradient boosted machines (GBM), support vector machines (SVM), and deep neural networks (DNNs)) in combination with two types of optimization techniques (random and Bayesian) and decide the k parameter of the k-fold cross validation. Prediction accuracy is defined as Pearson’s correlation coefficients (r) between empirical and predicted phenotype values, with model performance evaluated by how well models optimised on the training set perform on the test sets, as exemplified in Fig. 3b. The importance of individual features for model performance is also reported, except for SVR which use non-linear kernels and neural networks (Methods).

We evaluated four LASSO approaches for feature selection, finding them to perform equally well, and selecting similar numbers of important features (n=120-250), but with the LASSO grid being 2-3 orders of magnitudes faster than the alternatives (Supplementary Table 1). Irrespective of model type, the feature reduction from 8000 to ∼200 only marginally reduced model performance (Supplementary Fig. 6a and Supplementary Table 2). We further explored this marginal loss of model performance for the fastest method (ridge), finding the decrease to be most notable for CNVs (40% reduction), while hardly affecting proteomic and transcriptomic features (Supplementary Fig. 6b).

The reduction in features considerably (mean of 40 fold) reduced training times (Supplementary Fig. 6c), while resulting in a small loss of model performance for the training set (mean of 0.13) (Supplementary Fig. 6d). Because this small loss of model accuracy was somewhat greater in the training than for the test set, the reduction in features reduced overfitting, i.e. it improved the generalization of models from training to test sets. The Bayesian NN optimization failed to improve model predictions, and required careful fine-tuning of hyperparameters for each phenotype making it unsuitable for large-scale predictions. The other model permutation strategies in Gen-phen were benchmarked more in-depth using 30 representative phenotypes from the eight phenotype classes with LASSO grid feature selection (Fig. 3d). We found large variations in how well different phenotypes were predicted (Pearson’s r = 0.2 - 0.9), which likely reflects differences in the degree of environmental trait variation, but much lesser differences between model types. Overall, GBM Bayesian and Random regression models performed the best, giving the best predictions for ∼65% of the phenotypes, followed by SVR regression models (21 %) and neural networks (14%) (Fig. 3d). Overfitting was highest for the neural networks and lowest for SVR regression models (Supplementary Fig. 6e).

### Near perfect prediction accuracy using phenomics data

To determine which type of molecular data best predicts higher-level yeast phenotypes, we compared model performance using BayesGBM and different predictors as data inputs for the entire phenome. No significant difference between the distribution of accuracies among predictors was observed with BayesGBM, except for transcriptomic that consistently performed worse than other predictors (Fig. 4a-b). Across all model types with 30 test phenotypes previously defined, pangenome gene presence-absence data (PA) and P(AF) scores perform better, being superior predictors for an average (over model types) of 36% and 25% phenotypes respectively. LOF (15%) and proteomic data (15%) were least often the best predictors, and CNV (0.8%) almost never (Fig. 4c). When predicting higher-level phenotypes from the large SNP set (500K variants), training time increased drastically (from 5-10 min. to > 2 days per phenotype) and predictions were restricted to ∼20 random phenotypes (Supplementary Fig. 7a-b). BayesGBM did however not predict phenotypes from SNP data better than from other data types, hinting that the inclusion of genome-wide SNP data sets may not be necessary in machine learning predictions of higher-level phenotypes when other genomic data are available.

**Fig. 4.**
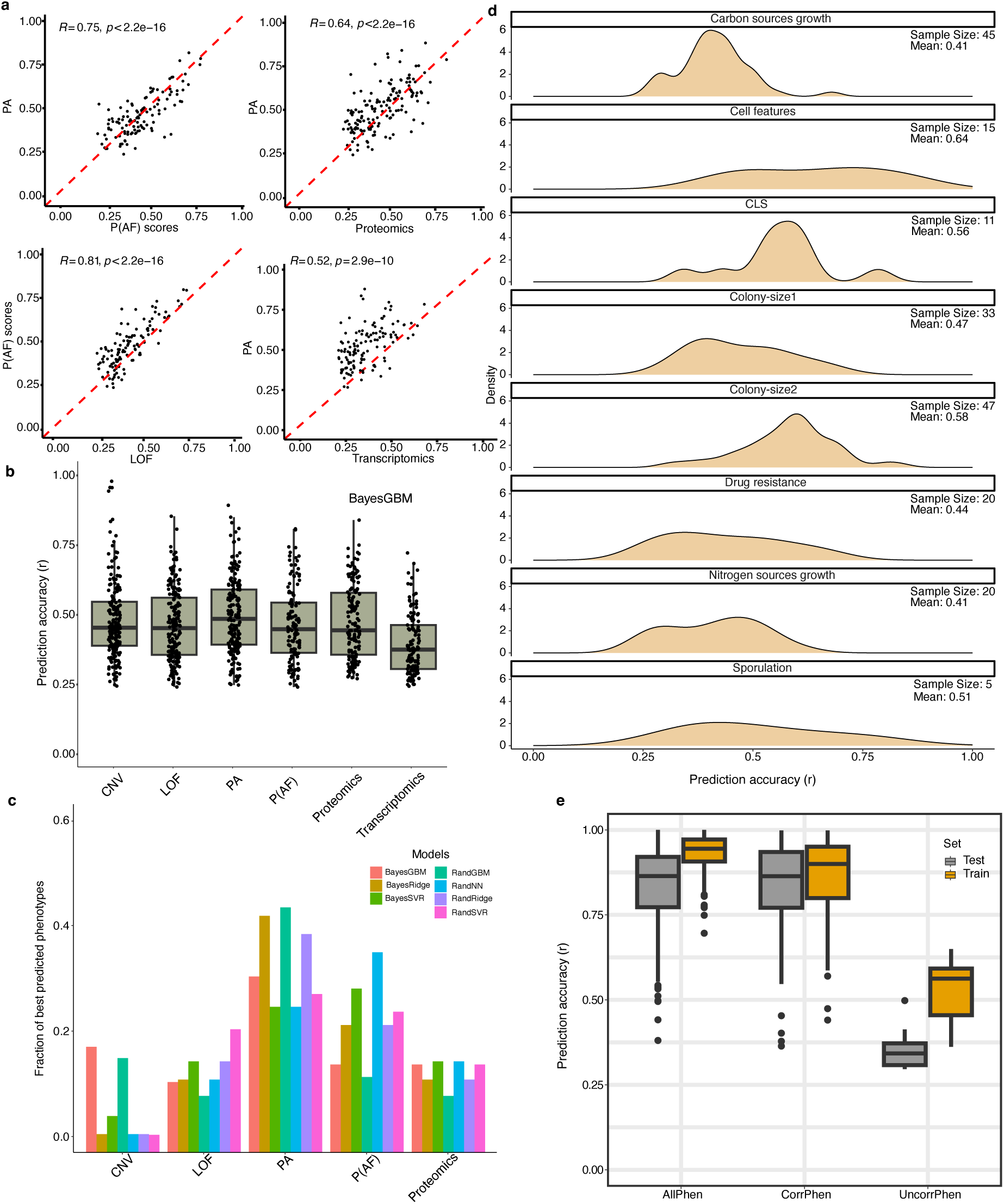
Phenotype prediction accuracies. **a,** Predictor comparisons based on prediction accuracies for the entire phenome. **b,** Prediction accuracies for the entire phenome based on different genomic and intermediate-level predictors. **c,** Fractions of phenotypes best predicted across predictors and methods for the 30 phenotypes test set. **d,** Comparing prediction accuracies for all phenotypes partitioned by classes show large variation. **e,** Predicting phenotypes using as predictors the entire phenome (AllPhen), correlated phenotypes (CorrPhen, with r >=0.3 or r <=-0.3) and uncorrelated phenotypes (UncorrPhen, −0.3<= r <=0.3).

Next, we selected the BayesGBM (as best model) and the pangenomic gene presence-absence data (as best predictor), to explore how well different phenotypes were predicted. Predicting all 223 phenotypes, we found larger variations in model performance, both between and within phenotypic classes (Fig. 4d). Overall, the best-predicted phenotype class was cell features (mean r = 0.64) followed by growth in stress conditions (colony-size 2, mean r = 0.58 and colony-size 1, mean r = 0.47), while growth under carbon and nitrogen source limitation was less well predicted (both 0.4).

We also used BayesGBM to investigate to what extent a single higher level phenotype can be predicted from a compendium of phenotypes, e.g. through effects of pleiotropic genes or genetic linkage across genome regions, but without explicit genomic predictor information. We tested three different scenarios, basing predictions of single higher level traits on all 223 such phenotypes, on only correlated phenotypes (r >0.3 or r< −0.3) and on only uncorrelated phenotypes (−0.3<r<0.3) respectively. Overall, using correlated phenotypes resulted in excellent predictions, with very high prediction accuracy (r >0.90) for 80 phenotypes and 32 showing near-perfect predictions (r >0.98) with a very small generalisation gap (difference between training and testing prediction accuracy, mean of 0.029, Fig. 4e). Only 10 phenotypes, all disconnected from other higher level phenotypes (Fig. 1b), were not well predicted (r<0.6). In contrast, including only uncorrelated phenotypes only rarely produced meaningful predictions (r<0.3 for 205 out of 223 phenotypes) and resulted in a bigger generalisation error (mean of 0.379, Fig. 4e). Finally, we explored whether incorporation of additional metadata in the form of strain ecological and geographic origin, heterozygosity level, phylogenetic clade and ploidy (1n, 2n, 4n) improved the accuracy of BayesGBM predictions from different data types, but found no meaningful improvements in any case (Supplementary Fig. 7c).

### Biological interpretation of machine-learning predictions

Next, we extracted the feature importance (FI) scores^32^, which quantify the contribution of individual features to the predictions, across all phenotypes and inspected whether the highest-scored features are enriched in some biological processes. The strength of the FI scores were independent of the frequency of the variant in the population, with most of the variants exhibiting low FI scores and only a few of them standing out (Fig. 5a-b). We observed few extremely high FI scores for phenotypes corresponding to growth measured in CuSO4 and Dyads production at 24h, with top candidates being YHR055C (*CUP1*) and YHR152W (*SPO12*) respectively (Fig. 5b). The gene *SPO12* was experimentally validated to determine dyads formation^17^, while the copy number variation of the *CUP1* gene is well known to be associated with yeast growth in CuSO4 environment^16^. Moreover, the LOF in *SPO12* is rare (present in <0.01 of the population) and therefore undetectable through GWAS. This suggests that ML can identify at least some rare genetic features of high impact associated with the phenotypes. For the cases where several genes were implicated in controlling the phenotype with moderate feature importance scores, we performed functional enrichment to investigate the underlying biological pathways. One such case is CLS CR day7 predicted using proteomics (Fig. 5a), the go-term analysis for the top genes (FI score>=0.01) showed functional enrichment in various processes including carbohydrate and glucose metabolic activities (Fig. 5c).

**Fig. 5.**
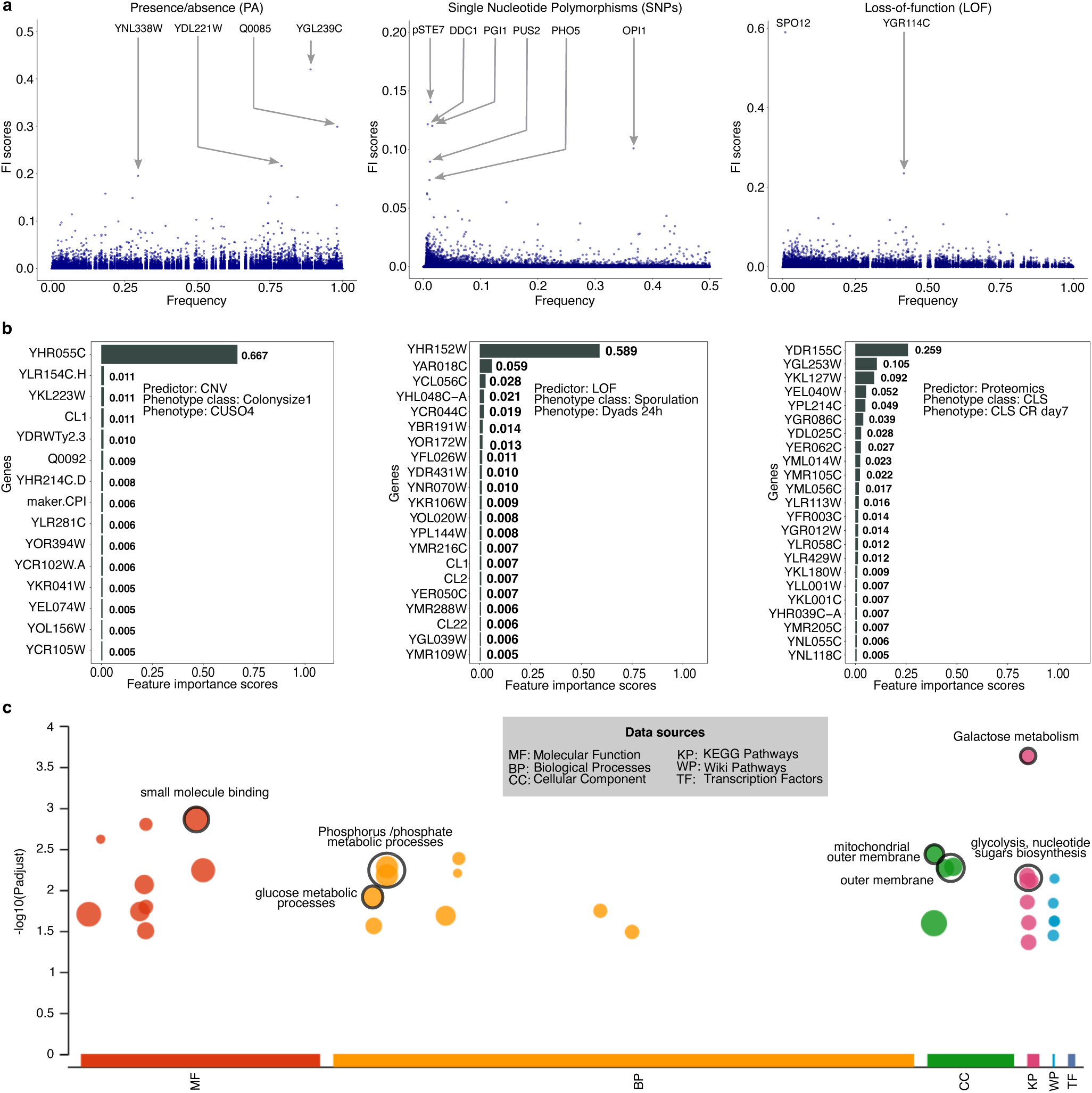
Biological interpretation of predictors. **a,** The strength of the FI scores for PA, SNPs, and LOF predictors is independent of their frequency in the population. The highest scoring variants are annotated. **b,** Top ranked feature importance score genes for growth yield in CuSO4, dyads production, and CLS CR day7. **c,** The enrichment analysis using the top scored genes from the prediction of CLS CR day7 using proteomics data.

FI scores distributions for most phenotypes were highly skewed, with most variants having very low average scores ranging from 0.002-0.0001 (0.002 /highest average for P(AF) scores and SNPs, Supplementary Fig. 9a-b). The number of features selected and the distribution of their scores varied significantly depending on the phenotypes. The phenotypes with highest average FI scores included growth measured in antifungal drugs for both P(AF) scores and SNPs (Supplementary Fig. 9a-b). Genes with high FI scores (> 0.01) for growth in tebuconazole and clotrimazole (highest average FI scores) showed enrichment in a number of processes, including hydrolase activity, siderophore transmembrane transport, transferase activity and nitrogen metabolic processes (Supplementary Fig. 9c-d).

## Discussion

We exploited the *S. cerevisiae* 1,011 framework to investigate the genotype-phenotype landscapes through machine learning. The availability of hundreds of phenotypes measured under controlled environments enabled us to explore phenotypic correlations at scale. We observed a strong impact of trait-type over environmental conditions in which the traits were recorded. Unexpected correlations between unrelated phenotypes also emerged, with several instances that were independently recapitulated using the gene-deletion collection^26^ demonstrating their biological basis, although only a modest overlap is expected between phenotypic consequences of natural variation and artificial deletions^33^. One interesting example is the correlation between the apparently unrelated CLS and azole resistance phenotypes, emerging from both the natural and the deletion collections. While the basis of such overlap is unknown, possible mechanism candidates include sterol lipids’ role in both membrane structures and signalling and genes involved in cell wall integrity^34–36^.

The genetic signals underlying phenotypic correlations can be captured by ML and thus improve predictions. Part of these signals are driven by pleiotropic variants that were also detected by GWAS, but some are driven by genetic determinants that escaped GWAS detection, such as epistatic interactions and rare variants. One unexpected finding of our study was the ability of ML to identify rare variants with high impact and their causative effects were supported by experimental validation. This points to ML being able to at least partially overcome one of the major limitations of GWAS. Rare variants, and rare epistatic interactions, are nevertheless likely to limit the power also of ML predictions in natural populations from individual genomes. Controlled crosses, where variants and variant pairs that are rare in natural populations are at much higher frequencies, can allow a more complete decomposition^37^ and accurate prediction^38^ of such traits. GWAS hits also included equal proportions of variants deriving from synonymous and missense mutations suggesting a non-neutral impact of synonymous mutations on complex phenotypes. Although the high frequency of synonymous mutation detected by GWAS could partially be derived from linkage disequilibrium, which limits the detection of the exact causal variants, their impact on complex traits remains an open question^39–42^. While several causative variants were previously validated, other SNPs including synonymous mutations, did not show detectable phenotypic effects when reconstructed in different backgrounds. This might be due to multiple factors, including our ability to detect small effects on phenotype from low impact variants and the genetic background dependency. The effect of yeast genetic background has been found to be strong also on gene deletion phenotypes^19^.

The construction of the Gen-Phen ML pipeline enabled us to systematically compare ML methods and input predictors. Pangenome consistently outperformed other predictors such as LOF, P(AF) scores, and omics data suggesting that the accessory genome plays an important role in deciding yeast phenotypes. These results mimic recent studies on bacteria antibiotic resistance, where pangenome variation is of even greater magnitude^43^. A limitation of our study is the number of samples available and increasing the sample size will certainly improve both the accuracy and the generalizability of the models. Combining different features did however not improve predictions. This might be due to the small sample size and the limitations of the ML methods used to model the complex interactions between different types of data and suggests the need for larger datasets along with multimodal ML methods that enable the integration of heterogeneous data^44–47^. Prediction accuracy among the test strains varies depending on their genetic distances from the training strains, with strains that are closely related better predicted than distantly related ones. Moreover, phenotypes measured in stress environments were predicted better than the phenotypes measured in nutrient-restricted environments, such as different types of carbon and nitrogen limitations. This was not unexpected since growth restricted by the availability of different carbon and nitrogen sources are high-complexity traits controlled by many enzymatic and regulatory genes and variants. Stress conditions are often simpler, with variants in a few genes responsible for excluding the stressor outside from cells having outsized phenotypic impact and being easier to detect^37,48^.

Overall, our study suggests that predicting complex traits based on genomic information in natural populations is feasible, but prediction accuracy varies greatly across phenotypes. Larger datasets can be further used to explore whether prediction accuracy improves and might enable us to identify common variants with small effects. Beyond the results presented, we provide a flexible ML workflow that can be easily adapted to different species and data types, including in clinical settings as an efficient way to predict phenotypes.

## Data availability

All data used in this study are available in the Supplementary Data file. The unpublished antifungal drug resistance data acquired within the CEFIPRA framework are available at: https://github.com/SakshiKhaiwal/CEFIPRA.

## Code availability

The developed computational pipeline and scripts are available at: https://github.com/SakshiKhaiwal/Genotype-to-phenotype-mapping-in-yeast/tree/main. This software is provided solely for the purpose of scientific reproduction of the research described in the associated article and general academic research. Any other use, including commercial use, is prohibited without prior written consent from the authors. All rights reserved.

## Acknowledgments

We thank Alan Moses, Marco Cosentino Lagomarsino, Marco Lorenzi, Yacine Ahmin, Ville Mustonen and Agnese Seminara for discussions and critical reading of the manuscript. This work was supported by Agence Nationale de la Recherche (ANR-11-LABX-0028-01, ANR-15-IDEX-01, ANR-18-CE12-0004, ANR-20-CE12-0020, ANR-22-CE12-0015), Fondation pour la Recherche Médicale (EQU202003010413), UCA AAP Start-up Deep tech, CEFIPRA, SATT Sud-Est to GL. This project received funding from the European Union’s Horizon 2020 research and innovation programme under grant agreement No 847581, the Region Sud, and the UCA J.E.D.I.

## Author contributions

S.K., M.D.C. and G.L. conceived the project and designed the experiments; S.K. developed the bioinformatic pipeline and performed all the primary analyses; S.S., B.P.B, J.W. designed and performed the phenotype experiments; I.B.H and P.B. contributed to the network expansion analysis; P.B., J.W. and G.L. contributed with resources and reagents; G.L. supervised the project; S.K. and G.L. wrote the paper with input from all the other authors.

## Material and Methods

### Datasets availability

#### Phenotypes for the natural population

The *S. cerevisiae* 1,011 collection has been phenotyped by multiple laboratories in different studies. We included 190 published phenotypes from four main publications^16–19^ and 33 unpublished phenotypes of which 22 phenotypes belong to growth measured in eight different antifungal drugs (https://github.com/SakshiKhaiwal/CEFIPRA) and 11 independently measured phenotypes, including cell size, DNA content, cell flocculation frequency, and CLS measured in rapamycin which are released in this study. Cell size and flocculation were estimated by flow cytometry. Each strain was grown in YPD media to exponential phase into 96-well microtiter plates prior to fixation with ethanol 70%. Cells were washed twice with PBS and resuspended in a DNA staining solution (15 µM propidium iodide (Sigma Aldrich), 100 µg/ml RNAse A (Sigma Aldrich), 0.1% Triton-X, in PBS solution) and incubated for 3 hours at 37°C in the dark. A total of 10’000 cells were directly analysed on a FACS Calibur flow cytometer using the FL2-A channel. The red fluorescence generated by stained DNA was used to identify isolated unbudded (1n) and budding (2n) cells, from which DNA content was estimated. Instead, cell size was inferred based on the FSC-H value of either unbudded and/or budding cells. The frequency of isolated 1n and 2n cells out of the total events was then leveraged to determine the extent of flocculation. Four replicates of the lab strain BY4743 were appended into each microtiter plate for data normalisation. The determination of CLS in rapamycin was performed in synthetic media supplemented with 0.025 µg/ml of drug as described here^49^. The phenotypes corresponding to doubling time have been inversed to represent the growth rates, so a higher value would mean faster growth and vice-versa. The phenome is enriched for the yield trait obtained from three main datasets^16,17,19^. In all the cases, yield is inferred from the size of the yeast colonies measured at a given time point. The entire set of phenotypes along with additional information corresponding to strains such as clades, ecological and geographical sources of isolation, etc. is given in Supplementary Data 2.

We divided the phenotypes into 8 major classes to make comparisons between various groups of environments, traits, and the experimental strategy (Supplementary Fig. 1a). For example, CLS, sporulation, cellular traits are distinguished from growth traits (yield/colony-size and growth rate). Growth rate and yield phenotypes which constitute 94 phenotypes measured in the same way (using scan-o-matic^50^) in three main types of environments, i.e. carbon-rich conditions, nitrogen-rich conditions, and antifungal drugs are separated into three classes namely ‘carbon sources growth’, ‘nitrogen sources growth’, and ‘drug resistance’ respectively. This was done to quantify the impact of the environment on the phenotypes. Finally, colony-sizes measured in similar stress conditions such as temperature, chemicals, and drugs constituting 35^16^ and 47^19^ phenotypes respectively, are obtained from two studies from different labs that allow us to compare the batch effects, experimental techniques, and timepoints for similar traits and environments. The phenotypes with defined classes with related publications are provided in Supplementary Data 3.

#### Gene-deletion collection Phenotypes

We extracted phenotypes from the gene deletion collection (yeastphenome.org), measured for similar traits and conditions of the ones available in the 1011 *S.cerevisiae* collection. We manually filtered the gene deletion collection first by traits (e.g. CLS, colony-size, sporulation) and then using keywords for environmental conditions. We obtained a list of 200 overlapping phenotypes (Supplementary Data 6-7).

#### Pangenome

Pangenomic matrices which include the matrix of presence/absence (PA) and copy number variations (CNV) of ORFs, loss of function (LOF) of the genes, and SNPs are obtained from^16^.

#### Population structure

The population structure is calculated using the same strategy as presented in^43^. The pairwise SNPs distance matrix was obtained from^16^ and the clustering is performed using the gengraph function of the adegenet package in R with the threshold range defined from 0.001 to the maximum value in the SNP distance matrix that varies the step size of 0.01 which gives us the range of clusters from 1 to 103.

#### P(AF) scores

The P(AF) scores^51^ that combine the functional effect of all variants in a gene were calculated for 1011 strains using the VariantAnnotation R package^52,53^.

#### Proteomics and transcriptomics

The proteomics and transcriptomics data are obtained from^54^ and^55^ respectively.

#### Phenotypic analysis

The correlation map is constructed using the corrplot 0.92 library in R 4.3.0. Phenotype networks are built using the correlation matrix using the network and ggnet2 functions of the network and ggplot2 library in R. We separated correlation and anticorrelation in distinct networks and filtered out weak correlations/anticorrelation using 0.5 and −0.3 thresholds respectively (Fig. 1b and Supplementary Fig. 1d). The phenomic distance between each pair of strains was calculated as the sum of the absolute differences of all the phenotypes and the tree was constructed using the biojn function from the ape library v5.7-1.

#### GWAS

We mined GWAS results from 99 phenotypes previously reported^17^. We re-run GWAS for 81 phenotypes using the same genetic data input and method to ensure consistency. Furthermore, we performed GWAS on 42 novel phenotypes that have not been included in any previous studies. GWAS are performed using similar settings and genome matrices (SNP, CNV, PA, and LOF)^16^. Genotype markers with < 5% MAF are removed from the analysis. Linear regression was implemented using fast linear mixture models using fast-lmm (fast-lmmc v2.07.20140723)^56^. The phenotypes are scaled to a standard normal distribution using the qqnorm function in R. We used the 5% family-wise error rate as threshold to extract significant markers related to a trait. This threshold for the p-value is obtained by performing 100 permutations of the phenotypes and extracting the fifth lowest p-value. The significant GWAS hits obtained from the GWAS analysis are annotated and for the SNPs that are not present in genes, we reported the flanking genes.

#### Linkage disequilibrium (LD) filtering

To remove any bias due to linkage disequilibrium (LD), we calculated LD scores for all pairs of SNPs that were within a 1kb window and less than 20 SNPs apart from each other. For pairs of SNPs with an LD score greater than 0.5 and significantly associated with the same phenotype, we only consider the SNP with a higher significance value (or lower p-value). LD scores are calculated using PLINK v1.90b6.18.

#### Variant effect predictions

The consequences of all the variants including the variants found to be significantly associated with specific phenotypes are calculated using the online web tool called Variant Effect Predictor (VEP) with a buffer size of 250 and upstream/downstream distance of 300 bp. To predict the impact of the missense variants to identify deleterious mutations, we used the mutfunc database and extracted all the predictions corresponding to the SNPs present in the pangenome of the *S. cerevisiae*^28^.

#### Network expansion analysis

Network expansion analysis is performed as described in^31^. To construct the interactome of the proteins, two sources of interactions namely STRING and BioGRID were used. To expand the signal, we used combined BioGRID and STRING with a score >=0.4. We used 173 phenotypes with at least 2 genes associated with an SNP (the bare minimum to start the network expansion). We ran the method and used the network propagation rank to measure the distance between the different traits in the study. To identify traits controlled by the same biological pathways, we extracted genetic clusters with more than 75% overlapping genes between different traits.

### Gen-phen prediction pipeline

The Gen-phen pipeline is built in Python 3.8 using the models from the scikit-learn 1.3.2 package. The pipeline comprises four major steps: pre-processing, feature selection, model learning, and model testing.

#### 1- Preprocessing

The Preprocessing step includes steps such as removing noisy data, imputing missing data, scaling the data, and splitting the data set into a training and test set. We have implemented three types of splitting criteria to split data into a training and a testing set, namely Hold-out at random (HOAR), Intra-clade hold-out (INHO), and Leave-one-clade-out (LOCO). HOAR involves randomly dividing genotypic and phenotypic data for all strains into a 75% training set and a 25% testing set, while INHO strategy only considers the strains belonging to the Wine European strains (approximately 300) and randomly assigns 50 of them to the test set and the rest to the training set. Finally, the LOCO strategy utilizes one entire clade as a test set, which in our case is ‘M3.Mosaic_Region_3’, and uses the rest of the clades for training. The data was scaled using standard scaling. The missing data in the features was imputed with the mean, while the samples missing the phenotypic data were removed from our analysis.

#### 2- Feature selection

LASSO regression is a commonly used machine learning technique that adds a penalty term to the regression function and sets the coefficient of some of the features to be zero which can be discarded from the data. The feature selection is implemented in two ways: 1) using the LASSO regression from scikit-learn and the high-dimensional LASSO regression using the Hi-LASSO Python library 1.0.6. Lasso selection is implemented through grid, random, and Bayesian search hyperparameter optimization. The grid and random hyperparameters optimization are done using GridSearchCV and RandomizedSearchCV function from scikit-learn while Bayesian hyperparameters optimization is done using BayesSearchCV function from scikit optimize 0.9.0. The grid selection is run with 5-fold cross-validation and with alphas equal to 0.001, 0.01, 0.1, and 0.5 for 10000 iterations. Random and Bayesian optimization is carried out using 5-fold cross-validation with alpha selected from a log-uniform distribution with values ranging from 1e-4 to 1 up to 500 and 200 iterations, respectively. The hi-lasso function is used with q1 and q2 being set to ‘auto’ with L and alpha being 30 and 0.01 respectively.

#### 3- Model learning

The model learning is implemented using 5 ML methods, namely ridge regression, elastic regression, support vector regression, gradient-boosted trees, and deep neural networks in combination with two types of hyperparameter optimization techniques, namely random and Bayesian search optimization along with five-fold cross-validation. All the hyperparameter distributions used for each model are defined in the ‘model’ part of the Gen-phen pipeline. Bayesian optimization is used with 100 iterations, whereas random optimization is used with 1000 iterations, except for the case of neural networks where we use 100 iterations due to their high computational power consumption.

#### Ridge Regression

Ridge regression is implemented using the Ridge function from the class linear models of the scikit-learn library and requires optimization of the hyperparameter, α which is a constant that controls the regularisation of the l2 norm. For randomised optimization, we used alpha from a log-uniform distribution with values ranging from 1 to 1000 while for Bayesian ridge regression, alpha is taken from a uniform distribution ranging from 1 to 10000.

#### Elastic Net Regression

Elastic net regression is also implemented using class linear models through the ElasticNet function from scikit-learn. It uses a combination of l1 and l2 and therefore requires optimization of two hyperparameters, α which is the constant that controls the penalty terms, and the l1 ratio known as the mixing parameter that controls the ratio of l1 and l2 regularisation. For both randomised and Bayesian optimization, we used uniform distributions with values ranging from 0.001 to 1 and from 0 to 1 for α and l1 ratio respectively.

#### Gradient Boosted Decision Trees

Gradient-boosted decision trees are ensemble-based methods and are implemented using the GradientBoostingRegressor function from the class ensemble in the scikit library. There are a number of hyperparameters that need to be defined for gradient-boosted trees. Some of them are the loss function, which is defined to be as ‘squared error’ in this case, ‘learning rate’ which is a parameter to control the contribution of each tree to the prediction, ‘n estimators’ to define the number of decision trees that are trained in the ensemble and so on.

#### Support Vector Regression

Support vector machines focus on trying to find a hyperplane that minimizes the error between the predicted and actual target values. This hyperplane could be a linear or non-linear function depending on the kernel used to map the data from a lower-dimensional to a higher-dimensional space for better clustering. Some of the hyperparameters to be optimised are ‘kernel’ for the type of kernel, the error margin ε is the range in which the loss function is equal to zero for data points that are predicted within it, regularization parameter ‘C’, and the kernel coefficient ‘gamma’. It is to be noted that the non-linear kernels in SVM transform the original feature space to a complex and non-linear higher-dimensional space, making the calculation of feature importance scores for those cases difficult.

#### Deep Neural Networks

The architecture of deep neural networks is defined mainly by two parameters, the number of hidden layers and the number of neurons in each hidden layer. A custom function is defined to design the neural network architecture depending on the dimension of the input layer and the percentage decrease in the number of nodes in each successive layer. This gives a neural network with a constant decrease in the number of nodes in the hidden layers. Some other hyperparameters that are optimized while training include α for the strength of the regularisation term, activation for hidden layers, solver for weight optimization, and batch size for mini-batches size for stochastic optimizers.

#### 4- Model testing

The accuracy of the model is assessed using the held-out test strains which consist of ∼150 strains that have not been used for training. The correlation between the predicted and experimentally measured values measured using Pearson’s coefficient is defined as the prediction accuracy. However, there are other variables such as r2 scores, and feature importance scores (wherever valid) can also be extracted for evaluating the model and features respectively.

#### Multivariate target predictions

We used three regression-based methods, multitask LASSO which is a linear-regression based method capable of building sparse models, a tree-based method namely multi-regression gradient boosting decision trees, and a neural network-based method that can be used to build highly non-linear and complex methods. To implement Multitarget LASSO, we used the ‘`linear_model.MultiTaskLasso’` function from the scikit-learn library along with grid-search hyperparameters optimization with three-fold cross-validation. We used the ‘`MultiOutputRegressor’` from ‘sklearn.multioutput’ along with the RandomizedSearchCV optimization to implement the multi-target gradient boosting decision trees regression. Finally, to construct a neural-network based multitarget regression model, we used a U-Net architecture with residual connections. This kind of architecture has the advantage of being easier to train, notably by avoiding the problem of vanishing gradients. This problem occurs particularly in deep neural networks, resulting in inefficient updating of the weights in the first layers^57^. The PyTorch library V1.13.1 with Python 3.8 was used for the construction of the neural net. The training, validation, and test sets were split randomly with 50%, 30%, and 20% strains respectively, followed by missing values imputation using the mean of the data for both features and target.

The number of neurons in the input and the output layers were set to be equal to the number of features and the number of phenotypes to be predicted, respectively. The model was built with eighteen hidden layers in addition to an input and output layer with three residual connections from the three layers at the beginning of the network to the three layers at the end of the network. We also used batch normalization which helps in increase the training efficiency and stability by recentering and rescaling the input layer. Furthermore, to regularise the training, dropout rates of 0.5, 0.25, and 0.25 were used for the beginning and end layers and middle layers respectively. Finally, a learning rate of 0.025 was used with the mean squared loss function and Adam optimizer (lr=0.0000001, weight_decay=0.1). Pearson’s coefficient between the predicted and the true values was used as a measure of accuracy and running accuracy is calculated as the average accuracy of the samples in the validation set. The training was performed for 200 epochs with validation being done once every five epochs.

The details of all the parameters and hyperparameters used for the individual phenotype prediction pipeline and multi-phenotype predictions can be found in the scripts at: https://github.com/SakshiKhaiwal/Genotype-to-phenotype-mapping-in-yeast/tree/main.

---

**Supplementary Note 1**

**Impact of population structure on prediction**

The *S. cerevisiae* has a fairly strong population structure and the collection is divided into 30 major clades^16^. This can represent a confounding factor and be treated as having real biological effects by ML models^58^. To reduce this bias, we measured the population structure and added it as an input feature for all predictions shown. Furthermore, to quantify the impact of the population, we tested the predictions derived from three splitting criteria: 1) Hold-out at random (HOAR), 2) Intra-clade hold out (INHO), and 3) LOCO (leave-one clade out). HOAR includes strains from all the clades (1,011 strains) and randomly divides them into a 75% training set and a 25% testing set. The INHO strategy considers only the Wine European clade (WE) with 276 strains which have a genome-wide SNP difference of only 0.12% (compared to an average of 0.5% overall) and a relatively weak internal population structure^16^, so both the training and testing samples come from almost an independent and identically distributed set. The strains are randomly divided into a testing set with 50 strains and the rest for training to maintain a sufficient number of strains. Finally, LOCO consists of using a complete clade (in this case, ‘M3.Mosaic_Region_3’) as the test clade and using the rest of the clades for training. In this case, the training and testing samples most likely have distributional differences arising due to the inherent biological structure. We performed the predictions for the 30 test phenotypes using ‘BayesGBM’ for all three strategies (Supplementary Fig. 6d). HOAR gave significant predictions for almost all phenotypes, while INHO gave significant predictions for only 50% of the phenotypes. However, the average accuracy in both cases remained at 0.5. The lower number of phenotypes predicted with INHO could be due to the significant decrease in the training samples (almost by 1/4th) compared to HOAR. The similar accuracies in the two cases imply no significant impact of the population structure while performing predictions at the population level. Moreover, using the LOCO strategy only 5 significant predictions were obtained with the same average accuracy of 0.5 as the above two cases. This suggests that predictions depend highly on the phenotypes and are predicted well if the relation between the genetic and phenotypic variation is generalized over the entire population (i.e. it is the same for all subclades). Additionally, we need to emphasize that the population size is one of the major constraints on the prediction accuracy, demonstrated by improved predictions with increased population size (Supplementary Fig. 7d). It is also to be noted that increasing training set size by adding technical replicates did not improve predictions further, signifying the necessity for larger biological populations for enhancing the prediction power.

**Supplementary Note 2**

**Using GWAS results to rank features and perform predictions**

To reduce the high dimensionality of input features, and to increase the efficiency and interpretability of ML models, it can be very important to perform feature selection in genomic predictions^59^. We used two main strategies to reduce the number of features for learning: 1) using multivariate LASSO regression strategies implemented in the Gen-phen pipeline (methods) to select a subset of features and 2) using a-prior knowledge from the GWAS results to rank genetic variants and selecting only the most relevant genetic markers. We ranked the genetic determinants using their association significance values (p-values) from GWAS and used the top 100, 200, and 500 markers as input features to predict the 30 test phenotypes (Supplementary Fig. 8). A comparison between the 8 methods shows that linear methods performed better in most cases when GWAS top markers were used as features (Supplementary Fig. 8a). In most cases, bayesian optimization-based models performed similar to the random hyperparameters optimization except for the neural networks where bayesian optimization consistently showed low performance. Comparing the number of markers used as features, we see small differences while increasing the number from 100 to 200 improving the performance in case of most phenotypes (Supplementary Fig. 8b). However, no significant differences were seen while increasing the number of markers to 500 from 200 and predictions improve slightly for some phenotypes while worsening for some others (Supplementary Fig. 8b). Furthermore, we compared the predictions among the different phenotype classes and see a very similar pattern as for the case when the pangenome was used as a predictor (Fig. 4c and Supplementary Fig. 8c). The best-predicted phenotypes included growth measured in stress environments with average accuracy ∼0.8 while the least predicted phenotypes included growth measured in nutrient-rich conditions (such as carbon and nitrogen). CLS, sporulation, and cell features were also predicted with relatively high accuracy (Pearson’s coefficient ∼0.7). In general, the predictions improved significantly when the a-priori information regarding the relevance of features was used compared to when no a-priori information was used and the entire pangenome was used as input (Fig. 4c and Supplementary Fig. 8c). However, we need to acknowledge that both the GWAS and the predictions were performed using the Euploid/Diploid strains which means both the training and the testing set had been included to perform the GWAS and this could potentially lead to an overestimation in the accuracy for the predictions using GWAS top markers. A bigger population size is required to detect this bias as further reducing the number of samples from GWAS can significantly reduce the robustness of the results.

**Supplementary Note 3**

**Multi-target regression models accurately predict the compound phenome**

To go beyond single-target predictions, we used two multi-target regression methods to construct models that can predict the 223 phenotypes at once (Supplementary Fig. 11a). We used the PA, P(AF) scores, and LOF as they were the best predictors across the 30 phenotypes test. We randomly divide data into a training (75%) and a testing (25%) set and run predictions using multitask LASSO, gradient-boosting decision trees, and a deep neural network. The overall average accuracy remained around 0.4 irrespective of the methods and input predictors (Supplementary Fig. 12). Despite being linear, Multitask LASSO performed comparable to the multi-regressor GBMs and deep neural network (Supplementary Fig. 12a-c). Moreover, the PA gave the best results among the three predictors (Supplementary Fig. 12a, d-e).

Next, we defined strain-level accuracy as all the phenotypes predicted vs. measured for each test strain and phenotype-level accuracy as all values predicted vs. measured for each phenotype considering all test strains. We observed significant variation in strain-level accuracy across clades, not depending on the clade’s size (Supplementary Fig. 11b and Supplementary Fig. 13a). Instead, accuracy per clade negatively correlated with the average genetic variation (number of ORFs and SNPs differences) per clade (Supplementary Fig. 14). This suggests the strains that are genetically similar to each other are predicted better compared to the distant strains. We also observed significant phenotype-level accuracy variation ranging from 0 to 0.75 (Supplementary Fig. 11c). Growth in nitrogen and sugar sources have the largest number of inaccurate predictions, consistent with results from individual phenotypes (Supplementary Fig. 13b). We extracted the feature importance for P(AF) scores as predictors from LASSO and GBMs. In both cases, *SFL1* was reported among the highest scored genes, which is also a top hit from the GWAS impacting many phenotypes. Furthermore, the top-scored genes from both cases are enriched in a number of important processes such as binding, cellular, and metabolic processes (Supplementary Data 21-22 and Supplementary Data 23-24).

**Supplementary Fig. 1.**
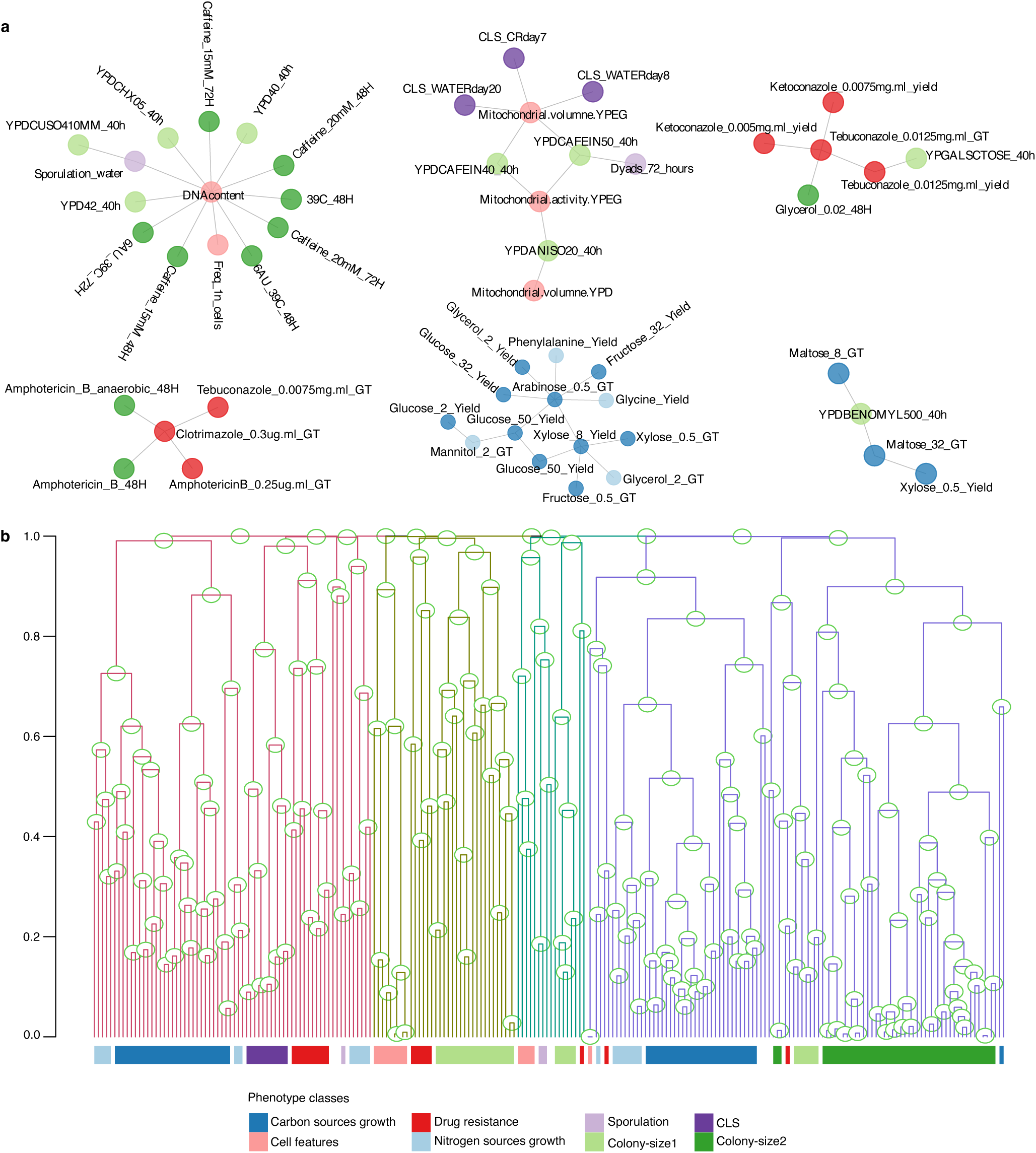
Phenotype anticorrelation networks and clustering. **a,** Anticorrelations produced six clusters (with Pearson’s coefficient <= −0.3) mainly consisting of trade-offs between growth rate and yield in various conditions. **b,** Hierarchical clustering groups phenotypes into four major groups (line colours) that are largely consistent with correlation patterns (Fig. 1a) and phenotypic classes (bottom coloured bar).

**Supplementary Fig. 2.**
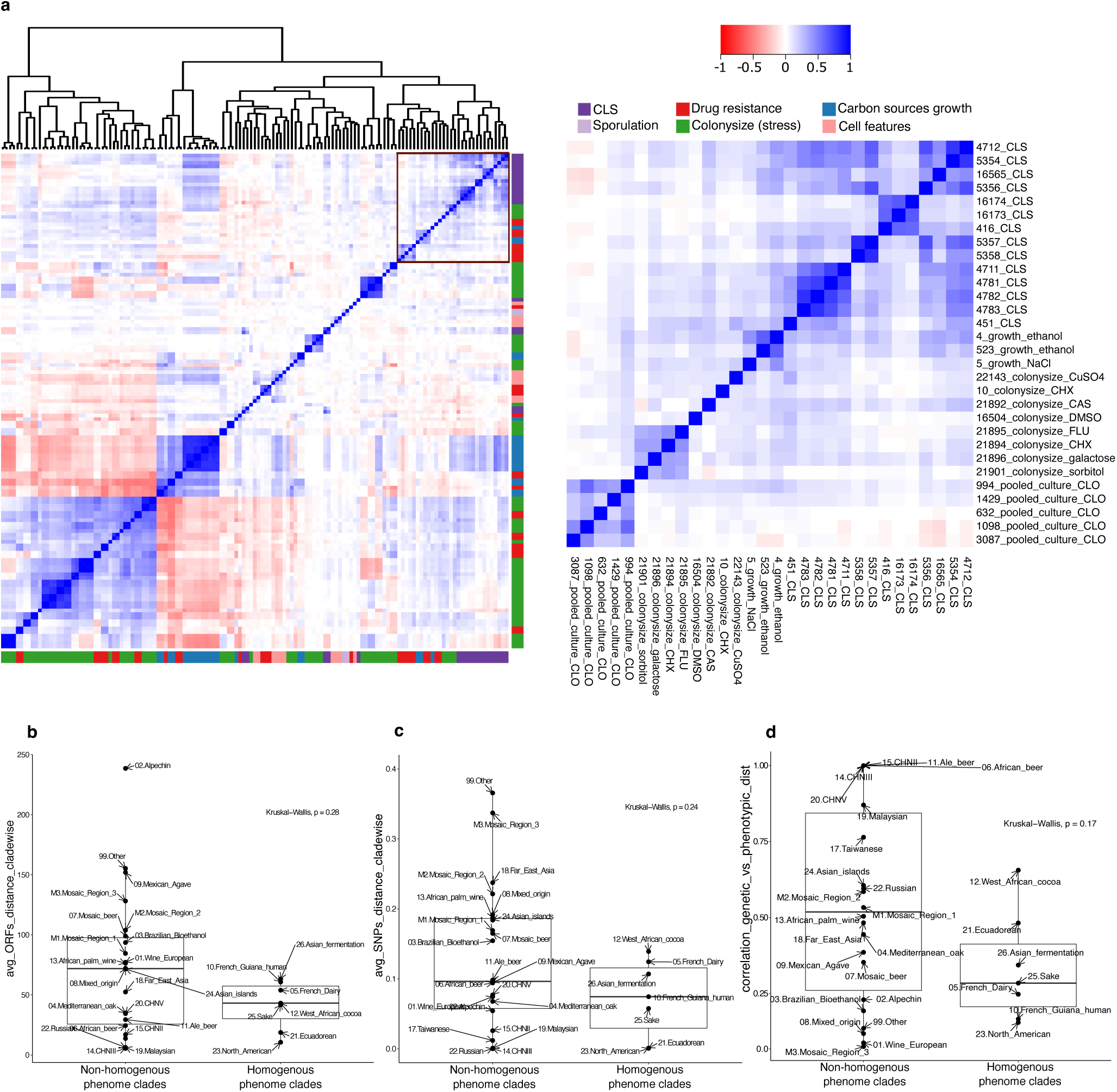
Phenotype correlations. **a,** Phenotype similarity matrix extracted for the gene-knockout phenotypes confirms similar patterns of correlations, including the antifungal drug resistance and CLS (right zoom-in). **b-d,** comparison between the average genetic variation between the clades highlighted in Fig. 1c and partitioned by homogenous (clades with strains clustered together based on their phenotypic distances) vs non-homogenous phenome (rest of the clades). **b,** The average SNPs difference between strains for each clade is lower for clades having a homogenous phenome’ compared to the other clades, except for clades with few strains. **c,** Average number of ORFs differences similarly showed lower variation for the clades with homogenous phenome. **d,** Correlation between genetic variation vs phenotypic variation also showed a similar trend.

**Supplementary Fig. 3.**
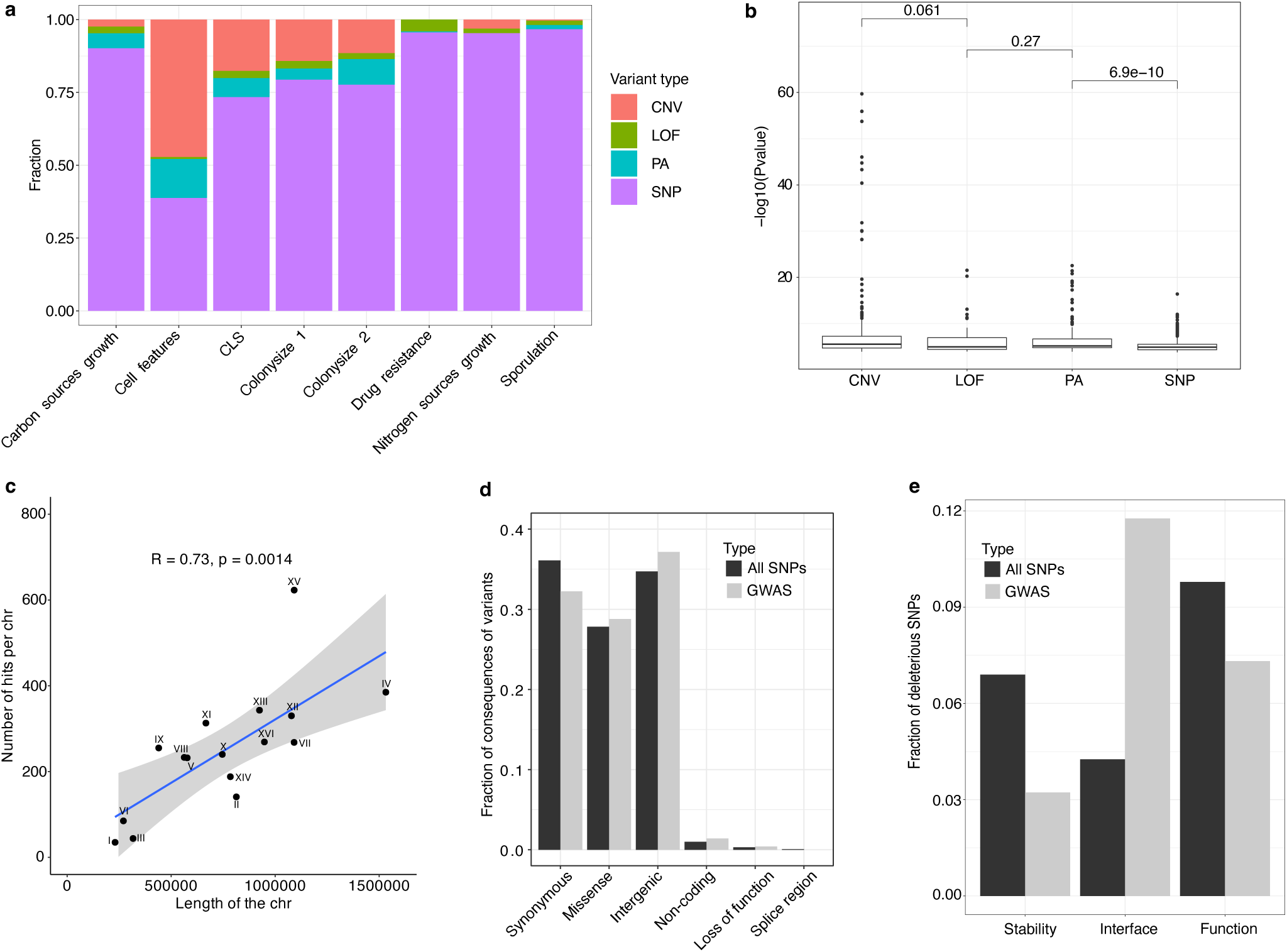
GWAS hits analysis and statistics. **a,** Fraction of various types of significant GWAS determinants in different phenotype classes. **b,** Comparison between the distributions of strength of associations between different types of genetic determinants (Wilcoxon test). **c,** The number of GWAS hits obtained for the entire phenome per chromosome shows a significant correlation with chromosome lengths, except for chromosome XV which has an exceptionally high number of hits. **d,** The number of variants in each category of the mutations obtained from the set of SNPs of the population reflects the proportion of variants obtained as hits from GWAS, except for variants in splice regions, however, this could be due to their small sample size (280 in the entire population). **e,** The fraction of deleterious SNPs in the SNPs set vs the GWAS hits shows that a higher number of SNPs impact the protein function compared to the stability of the protein. Although the number of deleterious SNPs affecting the protein-protein interface seems to be enriched in the GWAS, this is due to the very low number of predictions available for the interface interactions from mutfunc^28^ (Supplementary Data 11).

**Supplementary Fig. 4.**
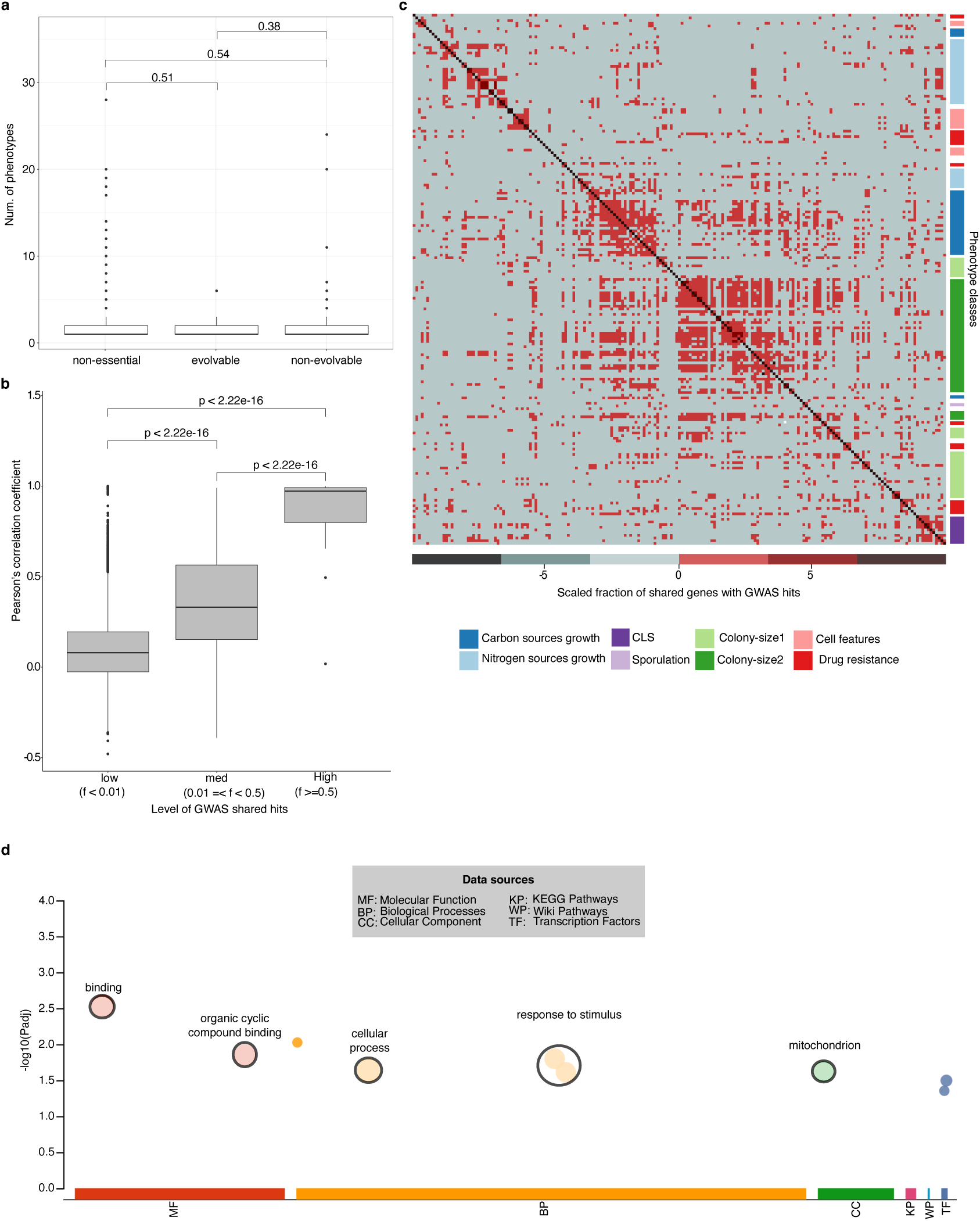
Pleiotropic genes with GWAS hits. **a,** The distributions of the number of phenotypes impacted per gene show no significant differences between the essential (evolvable and non-evolvable) and non-essential gene categories. **b,** We divided all pairs of phenotypes into three categories: low (<0.01), medium (0.01-0.5), and high (>0.5) fraction of shared GWAS hits. The fraction of GWAS hits shared between each pair of phenotypes is directly proportional to the level of correlation between the two. **c,** The number of shared genes with GWAS hits between each phenotype pair (heat map) reflects the pattern of the phenotypic correlation map (as visible from the right sidebar of the phenotypic classes). **d,** The pleiotropic genes (impacting more than 1 phenotype) from GWAS show enrichment in processes such as binding, organic cyclic compound binding, cellular response to stimulus, etc.

**Supplementary Fig. 5.**
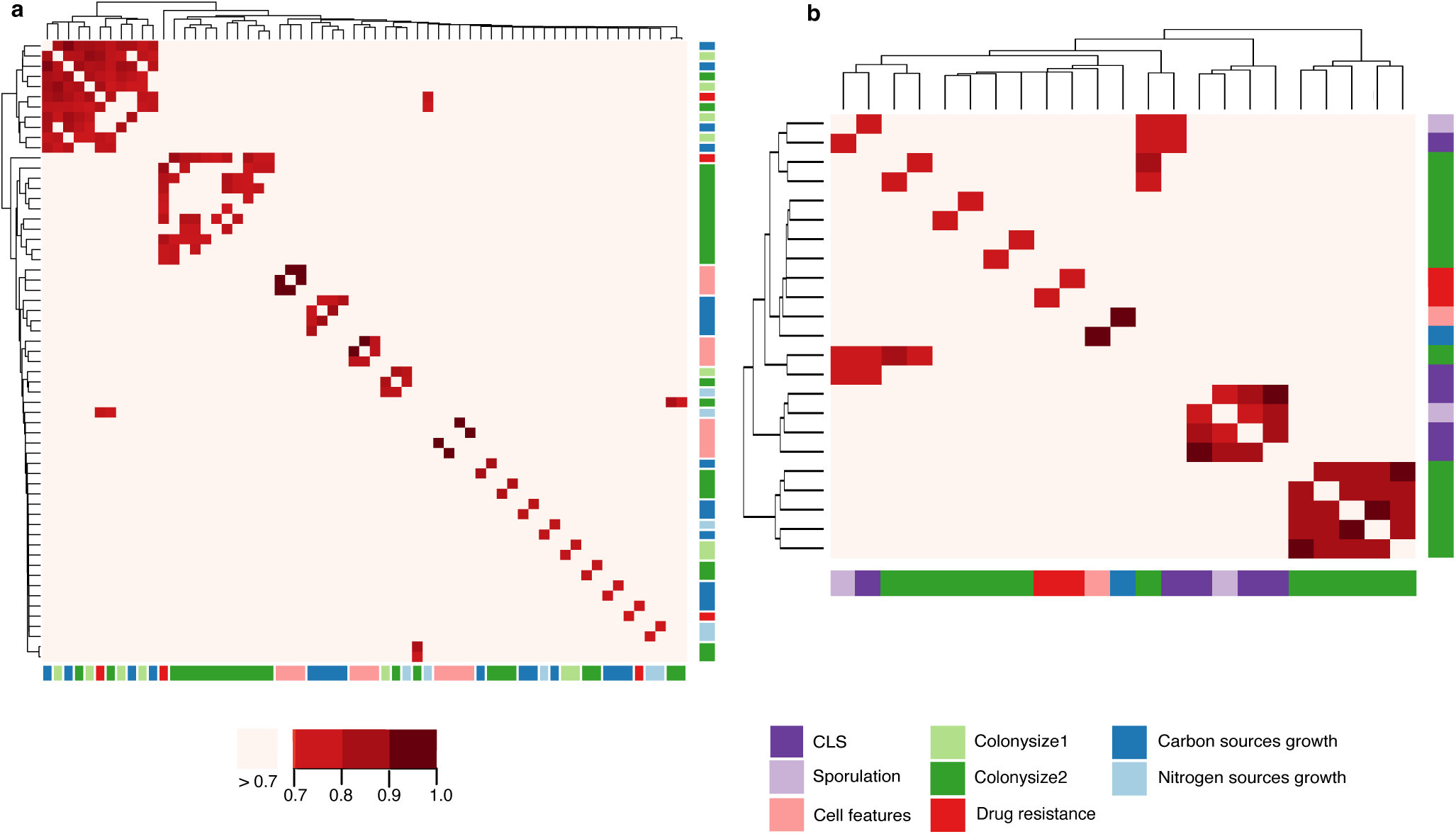
GWAS network expansion. **a-b** Network expansion analysis for the natural population and the gene-deletion collection respectively. The heatmap shows the percentage of overlapping genes between the gene modules associated with each phenotype. Only the phenotype pairs with 75% or higher number of overlapping genes are shown in red. **a,** Most phenotype pairs with a high number of overlapping modules are between phenotypes associated with similar conditions (along the diagonal) except the cluster on the top left position which consists of growth phenotypes measured in several carbon-rich (maltose, sorbitol, and galactose) and stress (caffeine, paraquat, and tebuconazole) conditions. **b,** Similar patterns emerge in the gene-deletion collection with most overlapping modules between similar traits and conditions. The two big clusters on the bottom right corner are among CLS and colony-size in stress conditions respectively.

**Supplementary Fig. 6.**
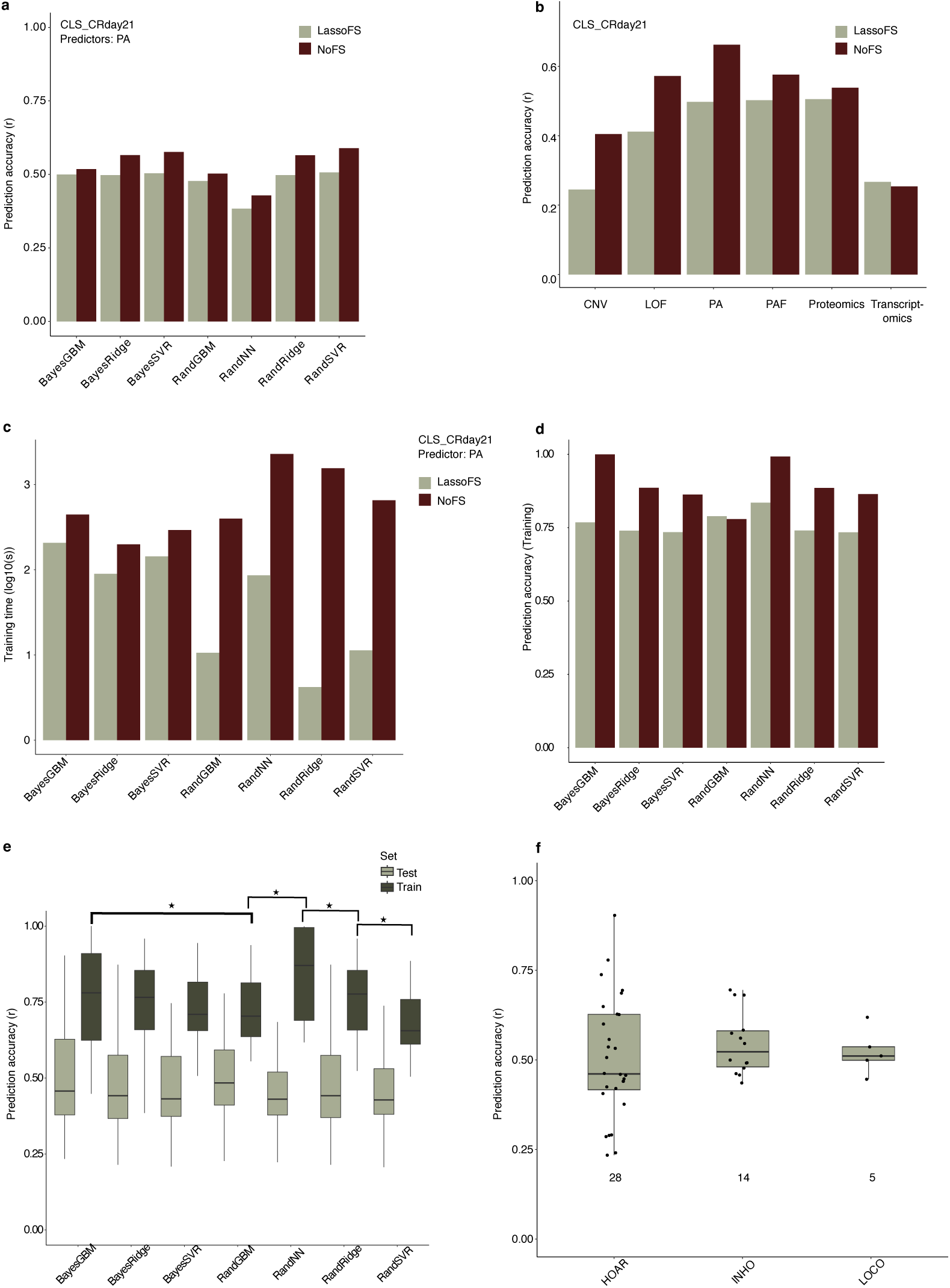
Feature selection and model benchmarking. **a,** Comparing prediction models with and without feature selection by LASSO regression **b,** Results were consistent across the different predictors tested using Ridge. **c,** The training times are reduced by a significant fraction for most of the methods when the number of features is reduced using an a-prior feature selection making the training process very efficient. **d,** The training accuracy was reduced for all methods when feature selection was performed. **e,** The difference between the training and the testing accuracy, generally known as the ‘generalization error’ is highest for neural networks and lowest for support vector regressors. **f,** The average prediction accuracy for the three splitting strategies i.e., hold-out at random (HOAR), intra-clade hold out (INHO) and leaving-one clade out (LOCO) tested for 30 phenotypes remained around 0.5. However, the number of phenotypes predicted reduced significantly from ∼95% for HOAR, to ∼0.45 for INHO and only ∼15% for LOCO.

**Supplementary Fig. 7.**
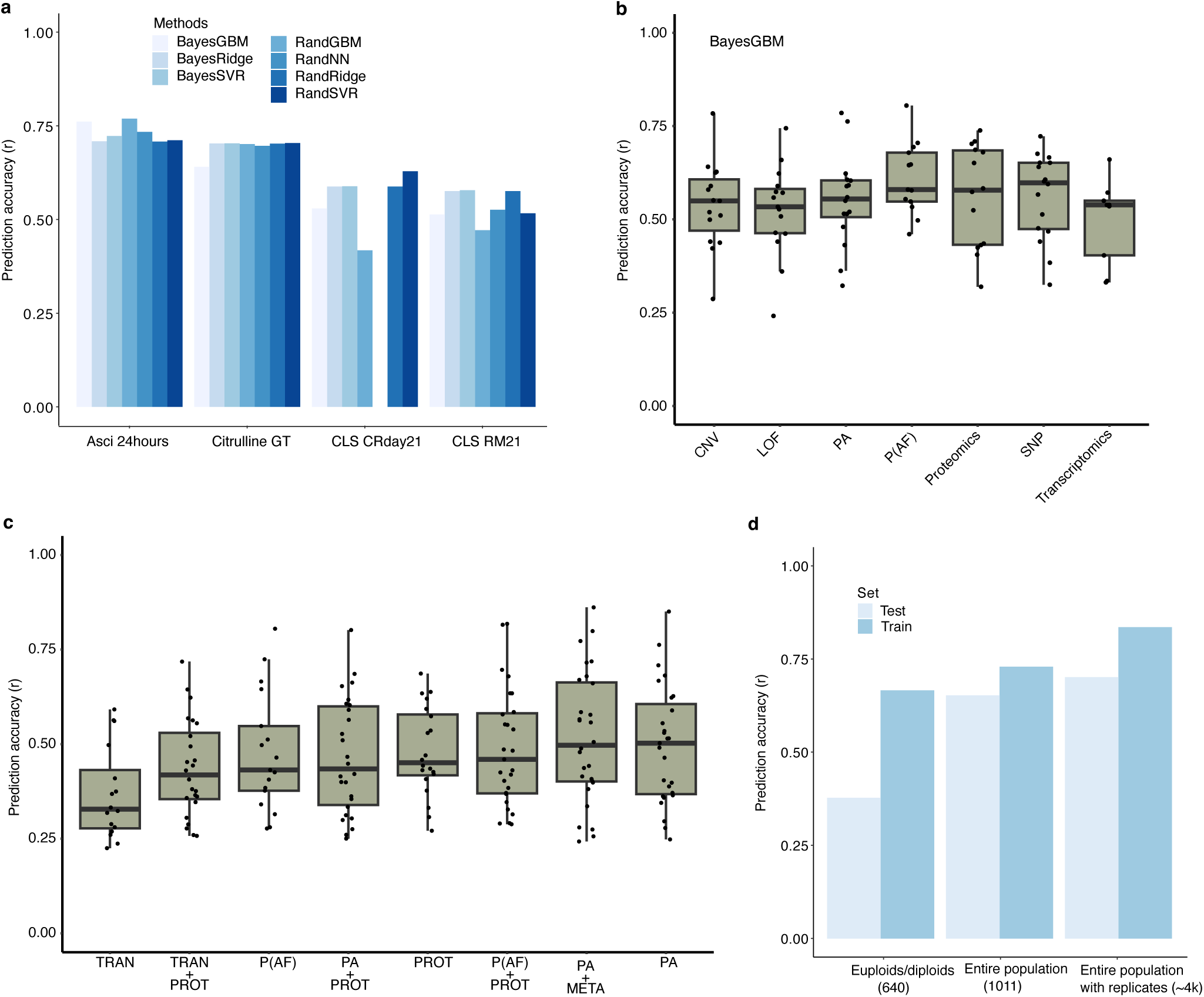
Benchmarking predictors. **a,** The phenotype predictions for four test phenotypes using SNPs (500k) as the predictor using all the seven methods show comparable results in most cases. **b,** The predictions for ∼20 random phenotypes are based on different predictors (PA, CNV, P(AF), LOF, SNP, transcriptomics, and proteomics). **c,** Comparison between different predictor combinations shows that the accuracy is rescued when combining proteomics with transcriptomics. **d,** The prediction accuracy increases while increasing the number of strains from 640 to 1011, however, remains comparable to the case when the technical replicates for each strain are considered distinct individuals. Moreover, the overfitting is reduced considerably with higher population sizes. The difference between the train and test set accuracy is negligible for the case when 1k and 4k strains were used compared to when only 650 strains were used.

**Supplementary Fig. 8.**
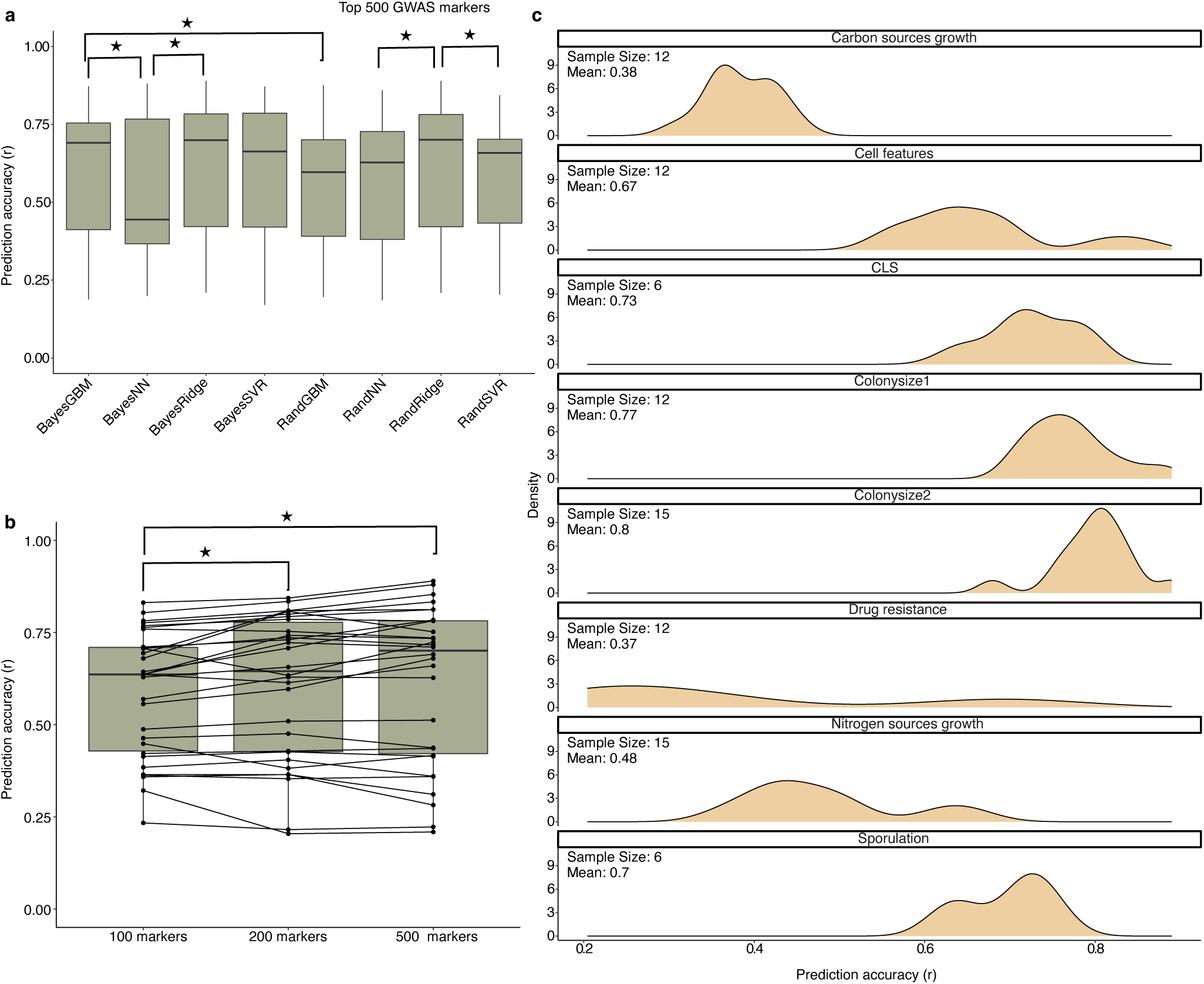
Top GWAS determinants as predictors. **a-c** show the prediction results using the top GWAS markers as predictors. **a**, The predictions using the top 500 GWAS markers are shown for the 30 test phenotypes for the eight methods. Linear-regression based methods such as ridge and SVR performed better than the non-linear methods such as GBM and neural networks for both random and Bayesian optimization. **b,** We compared the prediction accuracy with different numbers of input features, i.e. top 100, 200, and 500 GWAS markers. We see significant improvement in the accuracy when additional markers are added for most phenotypes, leading to an overall improvement in accuracy when considering a higher number of markers. **c,** Prediction accuracy for the entire set of phenotypes partitioned by classes. Growth measured in stress environments (colony-size1 and colony-size2) and CLS showed the highest average accuracy (∼0.8) while the classes with lowest predicted average accuracy (∼0.4) include growth measured in nutrient-rich conditions such as carbon and nutrient.

**Supplementary Fig. 9.**
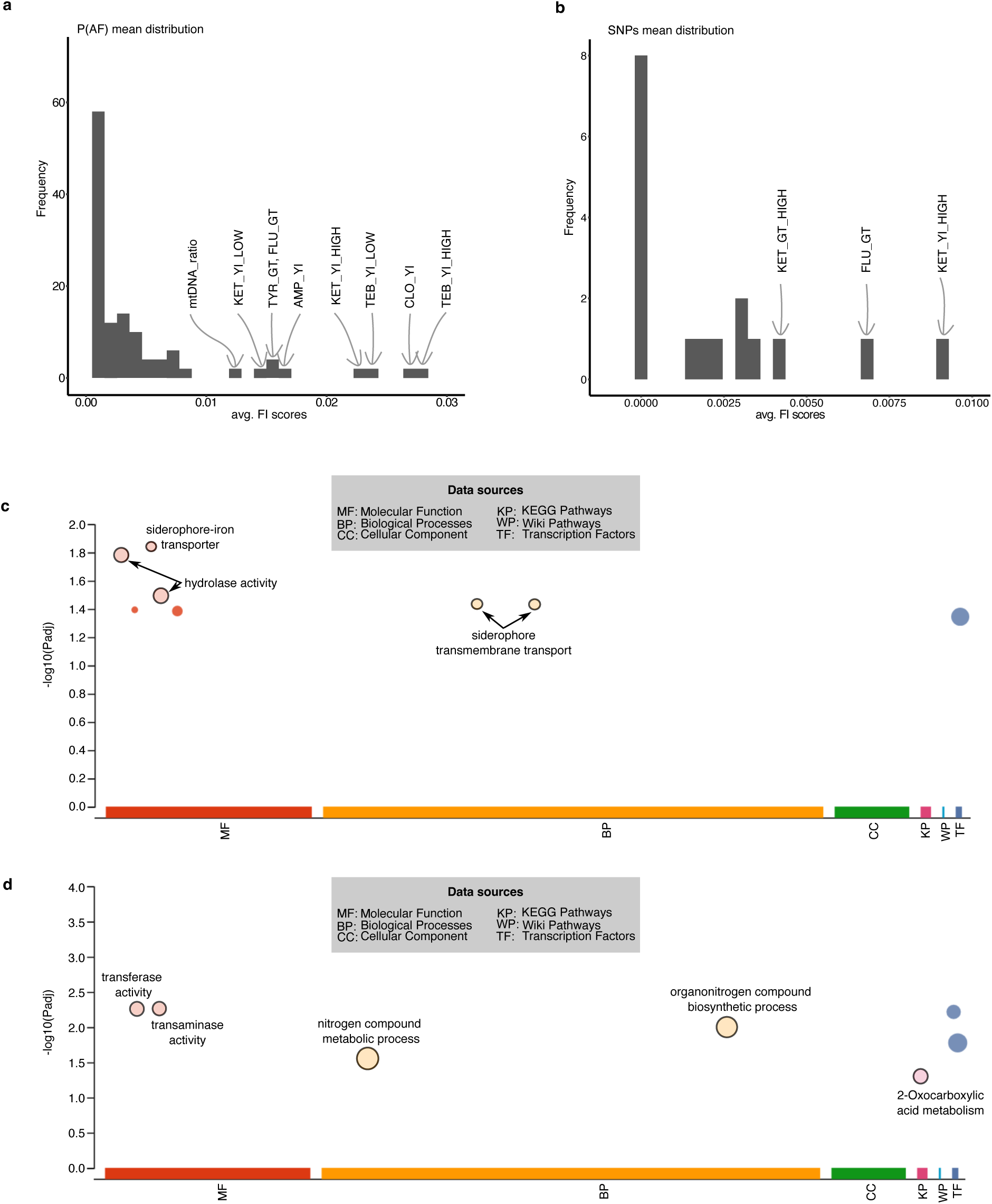
FI scores distributions and functional analysis of impactful features. Panels (a) and (b) show the distribution of average FI scores per phenotype using P(AF) scores and SNPs as the predictors respectively. **c-d** show respectively the go-enrichment results from gprofiler (https://biit.cs.ut.ee/gprofiler/gost) for the two top phenotypes, growth yield measured in tebuconazole and clotrimazole respectively. **c,** The genes with FI score > 0.01 for growth yield in tebuconazole show enrichment in various processes such as hydrolase activity, cyclin binding, etc. **d,** The genes with FI score > 0.01 for growth yield in clotrimazole show enrichment in transferase activity and nitrogen compound synthesis.

**Supplementary Figure 10.**
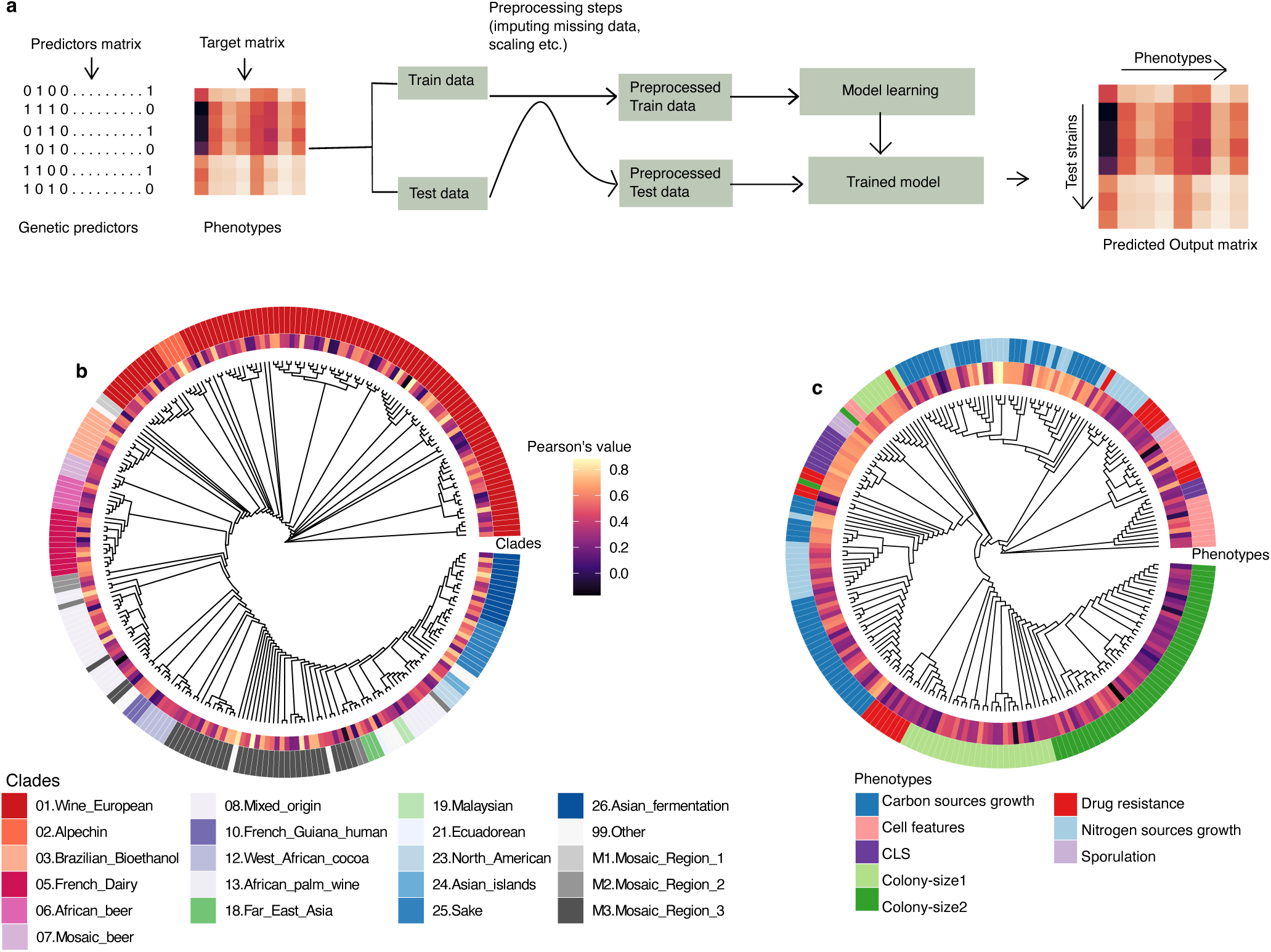
Multi-target prediction. **a,** Schematic of the multi-dimensional target depicting both the input predictor and output phenotype matrixes. The samples are represented as rows with features (PA, P(AF) scores, and LOF) and phenotypes as columns. **b,** Prediction accuracy (internal circle) across the phylogenetic tree using pangenome (PA) as the predictor and multitask LASSO as the model. **c,** Prediction accuracy (internal circle) among phenotype classes along the tree based on phenotypic correlations.

**Supplementary Fig. 11.**
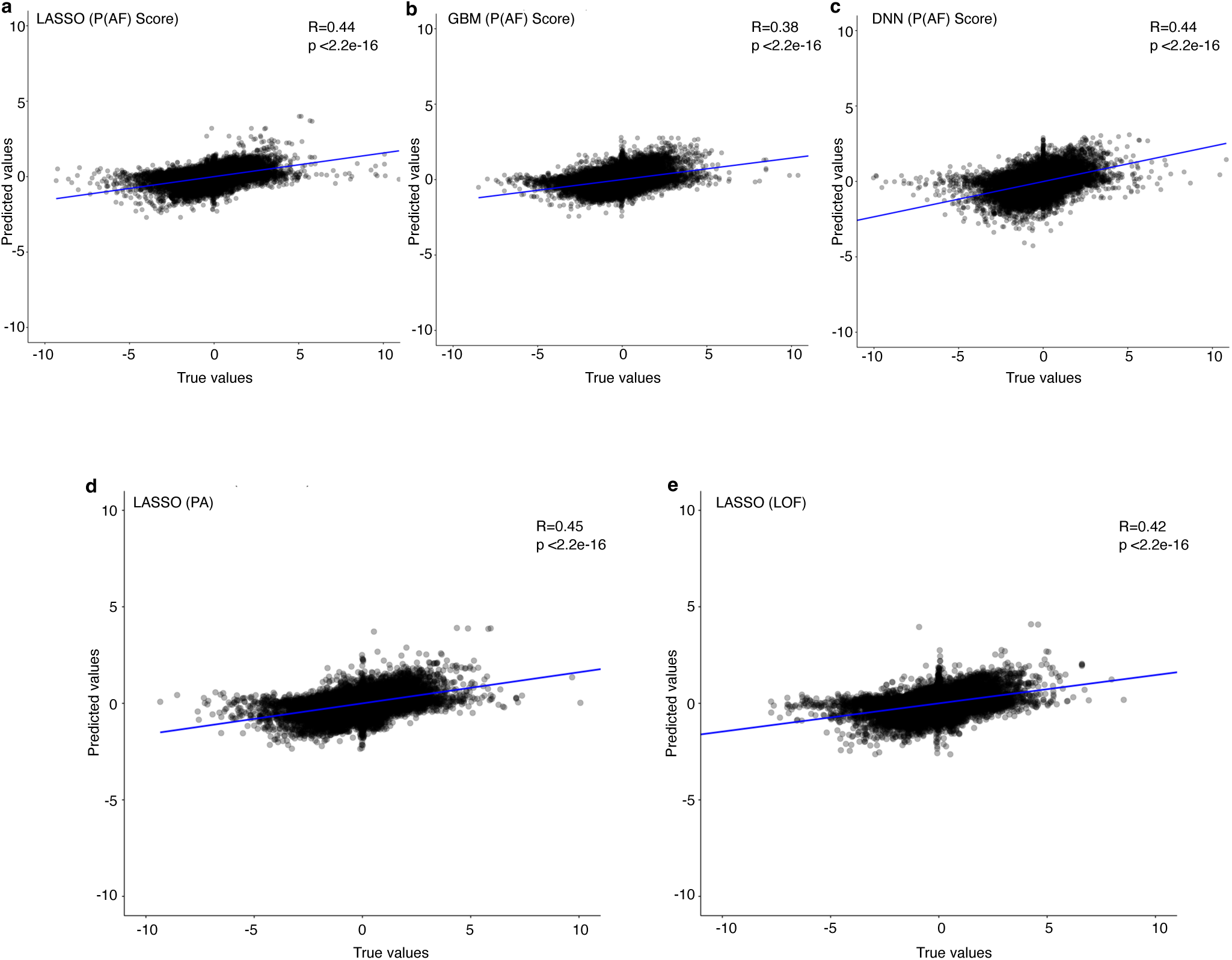
Entire phenome predictions. The prediction results measured as the Pearson’s correlation coefficient between the true and the predicted values for the entire phenome for the test set are shown for different methods and predictors. **a-c** prediction results for the entire phenome using different methods and (P(AF) scores) as input predictors. The prediction results from linear (LASSO) and non-linear (Neural networks) methods gave very similar results and were slightly better compared to the results from gradient boosting trees. **a, d-e** comparison between the prediction results while using different input predictors with multitask LASSO. All predictors gave similar predictions with Pearson’s coefficient around 0.4, however, predictions based on pangenome performed slightly better compared to the P(AF) scores and LOF.

**Supplementary Fig. 12.**
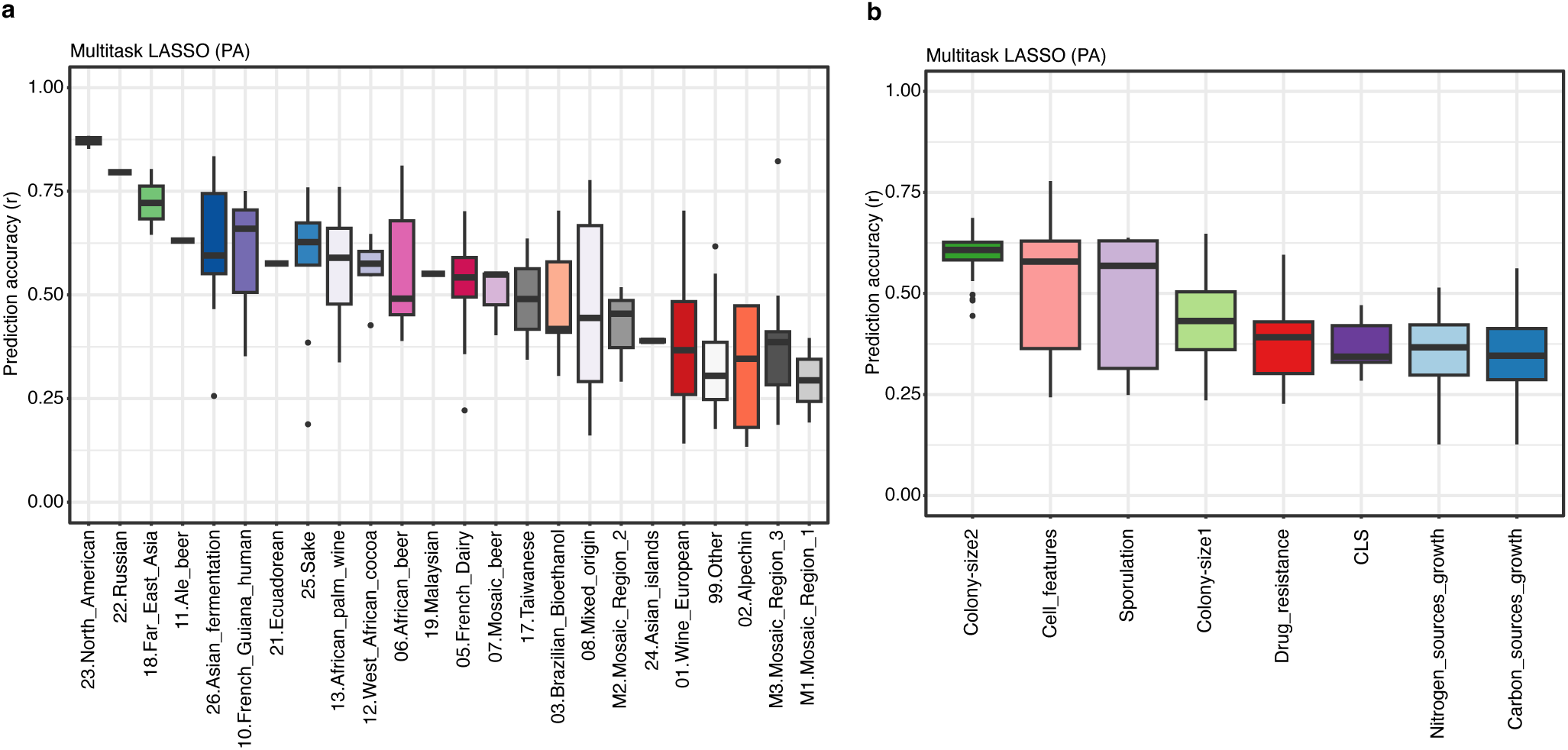
Phenome prediction by clades and phenotype classes. The prediction results for the entire phenome with pangenome and P(AF) scores as predictors and LASSO as prediction method are shown. **a,** The prediction accuracy for all phenotypes with test strains divided according to the clades they belong to shows a large variation ranging from 0.85 for North American strains to 0.3 for the strains belonging to mosaic clades. The accuracy remained independent of the size of the clades. **b,** The prediction results show significant variation among different phenotypic classes ranging from 0.6 for colony-size1 consisting of phenotypes measured in stress conditions to 0.4 for growth measured in nitrogen and carbon nutrients.

**Supplementary Fig. 13.**
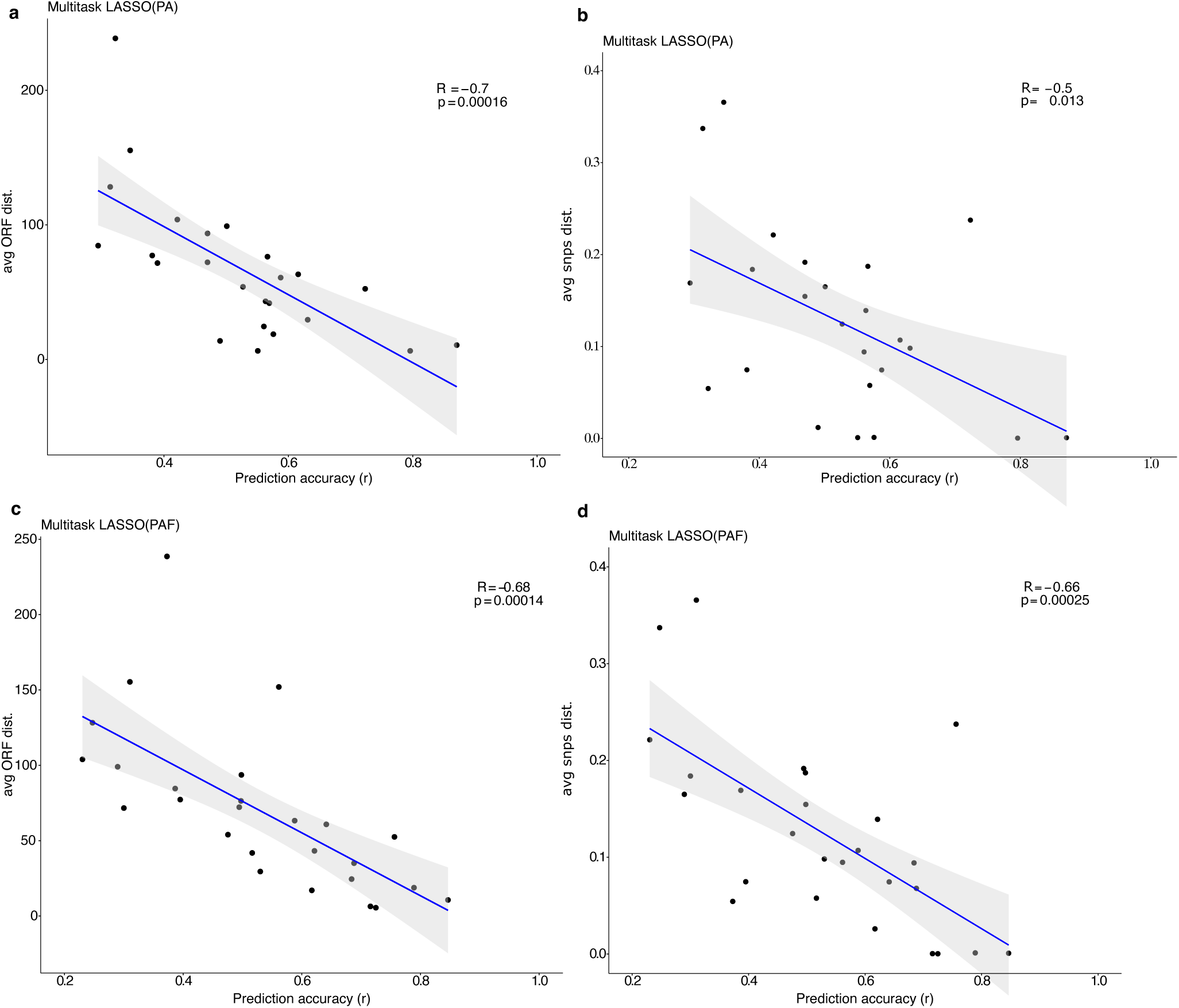
Prediction accuracy vs genetic distances. The average prediction accuracies per clade from multitask LASSO vs the average intra-clade genetic variation among the strains measured as the number of ORFs differences and the percentage of SNPs differences between each pair of differences using pangenome (panels (a) and (b)) and P(AF) scores (panels (c) and (d)) as predictors. In all cases, a significant anticorrelation is observed between the prediction accuracies and intra-clade genetic variation suggesting genetically closer strains phenome is easier to predict compared to distant strains.

**Supplementary Table 1.** Comparing different feature selection methods using CLS CR day 21 as the test phenotype and PA as test features shows similar accuracy and spareness (number of features selected). However, feature selection times vary greatly with LASSO grid being the most efficient method.

| Feature selection method | Time (s) | Accuracy | Num. features |
| --- | --- | --- | --- |
| Lasso grid | 7.7 | 0.52 | 204 |
| Lasso random | 767.9 | 0.49 | 250 |
| Lasso bayes | 794.1 | 0.57 | 120 |
| Hi-Lasso | 4444.6 | 0.52 | 212 |

**Supplementary Table 2.** Comparison between 4 ML methods and 2 optimization techniques for CLS CR day21 using PA show similar prediction accuracies, however a considerable difference between the training times. Random optimization was reported to be more efficient than Bayesian optimization. Moreover, Ridge and SVR were much more efficient that gradient boosting decision trees and neural networks. Bayesian neural networks required a lot of fine-tuning of the hyperparameters and high training time, however the prediction performance did not improve over other methods.

| Models | Bayesian optimization |  | Random optimization |  |
| --- | --- | --- | --- | --- |
|  | Train time (s) | Accuracy | Train time (s) | Accuracy |
| Ridge | 107.1 | 0.58 | 18.2 | 0.49 |
| GBM | 320.5 | 0.52 | 20.2 | 0.57 |
| SVR | 150.1 | 0.45 | 29.5 | 0.57 |
| Neural networks | 415.6 | 0.46 | 100.9 | 0.52 |

## References

1. Goddard, M. E., Kemper, K. E., MacLeod, I. M., Chamberlain, A. J. & Hayes, B. J. Genetics of complex traits: prediction of phenotype, identification of causal polymorphisms and genetic architecture. Proc. Biol. Sci. 283, 20160569 (2016).

2. Ehrenreich, I. M. et al. Dissection of genetically complex traits with extremely large pools of yeast segregants. Nature 464, 1039–1042 (2010).

3. Zhang, H. et al. Genome-wide association study identifies 32 novel breast cancer susceptibility loci from overall and subtype-specific analyses. Nat. Genet. 52, 572–581 (2020).

4. de Lange, K. M. et al. Genome-wide association study implicates immune activation of multiple integrin genes in inflammatory bowel disease. Nat. Genet. 49, 256–261 (2017).

5. Li, Z. et al. Genome-wide association analysis identifies 30 new susceptibility loci for schizophrenia. Nat. Genet. 49, 1576–1583 (2017).

6. Tian, D. et al. GWAS Atlas: a curated resource of genome-wide variant-trait associations in plants and animals. Nucleic Acids Res. 48, D927–D932 (2020).

7. Tam, V. et al. Benefits and limitations of genome-wide association studies. Nat. Rev. Genet. 20, 467–484 (2019).

8. Guo, T. & Li, X. Machine learning for predicting phenotype from genotype and environment. Curr. Opin. Biotechnol. 79, 102853 (2023).

9. Lakiotaki, K. et al. Automated machine learning for genome wide association studies. Bioinformatics 39, btad545 (2023).

10. Schrag, T. A. et al. Beyond Genomic Prediction: Combining Different Types of omics Data Can Improve Prediction of Hybrid Performance in Maize. Genetics 208, 1373–1385 (2018).

11. Cheng, C.-Y. et al. Evolutionarily informed machine learning enhances the power of predictive gene-to-phenotype relationships. Nat. Commun. 12, 5627 (2021).

12. Yeh, C.-L. C., Jiang, P. & Dunham, M. J. High-throughput approaches to functional characterization of genetic variation in yeast. Curr. Opin. Genet. Dev. 76, 101979 (2022).

13. Liti, G., Warringer, J. & Blomberg, A. Mapping Quantitative Trait Loci in Yeast. Cold Spring Harb. Protoc. 2017, pdb.prot089060 (2017).

14. Fay, J. C. The molecular basis of phenotypic variation in yeast. Curr. Opin. Genet. Dev. 23, 672–677 (2013).

15. Liti, G. & Louis, E. J. Advances in Quantitative Trait Analysis in Yeast. PLoS Genet. 8, e1002912 (2012).

16. Peter, J. et al. Genome evolution across 1,011 Saccharomyces cerevisiae isolates. Nature 556, 339–344 (2018).

17. De Chiara, M. et al. Domestication reprogrammed the budding yeast life cycle. Nat. Ecol. Evol. 6, 448–460 (2022).

18. D’Angiolo, M. et al. Telomeres are shorter in wild Saccharomyces cerevisiae isolates than in domesticated ones. Genetics 223, iyac186 (2023).

19. Galardini, M. et al. The impact of the genetic background on gene deletion phenotypes in Saccharomyces cerevisiae. Mol. Syst. Biol. 15, e8831 (2019).

20. Caudal, É. et al. Pan-transcriptome reveals a large accessory genome contribution to gene expression variation in yeast. Nat. Genet. 56, 1278–1287 (2024).

21. Muenzner, J. et al. Natural proteome diversity links aneuploidy tolerance to protein turnover. Nature 630, 149–157 (2024).

22. Sharon, E. et al. Functional Genetic Variants Revealed by Massively Parallel Precise Genome Editing. Cell 175, 544–557.e16 (2018).

23. Roy, K. R. et al. Multiplexed precision genome editing with trackable genomic barcodes in yeast. Nat. Biotechnol. 36, 512–520 (2018).

24. Draskau, M. K. & Svingen, T. Azole Fungicides and Their Endocrine Disrupting Properties: Perspectives on Sex Hormone-Dependent Reproductive Development. Front. Toxicol. 4, 883254 (2022).

25. Wei, X. & Zhang, J. Environment-dependent pleiotropic effects of mutations on the maximum growth rate r and carrying capacity K of population growth. PLoS Biol. 17, e3000121 (2019).

26. Turco, G. et al. Global analysis of the yeast knockout phenome. Sci. Adv. 9, eadg5702 (2023).

27. Baccarini, L., Martínez-Montañés, F., Rossi, S., Proft, M. & Portela, P. PKA-chromatin association at stress responsive target genes from Saccharomyces cerevisiae. Biochim. Biophys. Acta 1849, 1329–1339 (2015).

28. Wagih, O. et al. A resource of variant effect predictions of single nucleotide variants in model organisms. Mol. Syst. Biol. 14, e8430 (2018).

29. Hu, X. et al. Structural and mechanistic insights into fungal β-1,3-glucan synthase FKS1. Nature 616, 190–198 (2023).

30. Cowen, L., Ideker, T., Raphael, B. J. & Sharan, R. Network propagation: a universal amplifier of genetic associations. Nat. Rev. Genet. 18, 551–562 (2017).

31. Barrio-Hernandez, I. et al. Network expansion of genetic associations defines a pleiotropy map of human cell biology. Nat. Genet. 55, 389–398 (2023).

32. Huynh-Thu, V. A., Saeys, Y., Wehenkel, L. & Geurts, P. Statistical interpretation of machine learning-based feature importance scores for biomarker discovery. Bioinformatics 28, 1766–1774 (2012).

33. Jiang, D. & Zhang, J. Detecting natural selection in trait-trait coevolution. BMC Ecol. Evol. 23, 50 (2023).

34. Matecic, M. et al. A microarray-based genetic screen for yeast chronological aging factors. PLoS Genet. 6, e1000921 (2010).

35. Garay, E. et al. High-resolution profiling of stationary-phase survival reveals yeast longevity factors and their genetic interactions. PLoS Genet. 10, e1004168 (2014).

36. Anderson, J. B. et al. Mode of selection and experimental evolution of antifungal drug resistance in Saccharomyces cerevisiae. Genetics 163, 1287–1298 (2003).

37. Hallin, J. et al. Powerful decomposition of complex traits in a diploid model. Nat. Commun. 7, 13311 (2016).

38. Märtens, K., Hallin, J., Warringer, J., Liti, G. & Parts, L. Predicting quantitative traits from genome and phenome with near perfect accuracy. Nat. Commun. 7, 11512 (2016).

39. Hunt, R. C., Simhadri, V. L., Iandoli, M., Sauna, Z. E. & Kimchi-Sarfaty, C. Exposing synonymous mutations. Trends Genet. TIG 30, 308–321 (2014).

40. Bailey, S. F., Alonso Morales, L. A. & Kassen, R. Effects of Synonymous Mutations beyond Codon Bias: The Evidence for Adaptive Synonymous Substitutions from Microbial Evolution Experiments. Genome Biol. Evol. 13, evab141 (2021).

41. Shen, X., Song, S., Li, C. & Zhang, J. Synonymous mutations in representative yeast genes are mostly strongly non-neutral. Nature 606, 725–731 (2022).

42. She, R. & Jarosz, D. F. Mapping Causal Variants with Single-Nucleotide Resolution Reveals Biochemical Drivers of Phenotypic Change. Cell 172, 478–490.e15 (2018).

43. Moradigaravand, D. et al. Prediction of antibiotic resistance in Escherichia coli from large-scale pan-genome data. PLOS Comput. Biol. 14, e1006258 (2018).

44. Argelaguet, R. et al. Multi-Omics Factor Analysis—a framework for unsupervised integration of multi-omics data sets. Mol. Syst. Biol. 14, e8124 (2018).

45. Tan, X., Su, A. T., Hajiabadi, H., Tran, M. & Nguyen, Q. Applying Machine Learning for Integration of Multi-Modal Genomics Data and Imaging Data to Quantify Heterogeneity in Tumour Tissues. Methods Mol. Biol. Clifton NJ 2190, 209–228 (2021).

46. Parisot, S. et al. Disease prediction using graph convolutional networks: Application to Autism Spectrum Disorder and Alzheimer’s disease. Med. Image Anal. 48, 117–130 (2018).

47. Nguyen, N. D., Huang, J. & Wang, D. A deep manifold-regularized learning model for improving phenotype prediction from multi-modal data. Nat. Comput. Sci. 2, 38–46 (2022).

48. Vázquez-García, I. et al. Clonal Heterogeneity Influences the Fate of New Adaptive Mutations. Cell Rep. 21, 732–744 (2017).

49. Barré, B. P. et al. Intragenic repeat expansion in the cell wall protein gene HPF1 controls yeast chronological aging. Genome Res. 30, 697–710 (2020).

50. Zackrisson, M. et al. Scan-o-matic: High-Resolution Microbial Phenomics at a Massive Scale. G3 GenesGenomesGenetics 6, 3003–3014 (2016).

51. Jelier, R., Semple, J. I., Garcia-Verdugo, R. & Lehner, B. Predicting phenotypic variation in yeast from individual genome sequences. Nat. Genet. 43, 1270–1274 (2011).

52. Obenchain, V. et al. VariantAnnotation : a Bioconductor package for exploration and annotation of genetic variants. Bioinformatics 30, 2076–2078 (2014).

53. Dunham, A. Modelling the structural, functional and phenotypic consequences of protein coding mutations. ([object Object], 2021). doi:10.17863/CAM.82589.

54. Muenzner, J. et al. The natural diversity of the yeast proteome reveals chromosome-wide dosage compensation in aneuploids. Preprint at 10.1101/2022.04.06.487392 (2022).

55. Caudal, E. et al. Pan-transcriptome reveals a large accessory genome contribution to gene expression variation in yeast. Preprint at 10.1101/2023.05.17.541122 (2023).

56. Lippert, C. et al. FaST linear mixed models for genome-wide association studies. Nat. Methods 8, 833–835 (2011).

57. He, K., Zhang, X., Ren, S. & Sun, J. Deep Residual Learning for Image Recognition. (2015) doi:10.48550/ARXIV.1512.03385.

58. Whalen, S., Schreiber, J., Noble, W. S. & Pollard, K. S. Navigating the pitfalls of applying machine learning in genomics. Nat. Rev. Genet. 23, 169–181 (2022).

59. Libbrecht, M. W. & Noble, W. S. Machine learning applications in genetics and genomics. Nat. Rev. Genet. 16, 321–332 (2015).

